# CRISPR Screening in Tandem with Targeted mtDNA Damage Reveals WRNIP1 Essentiality

**DOI:** 10.1101/2023.10.03.560559

**Authors:** Tanja Sack, Piriththiv Dhavarasa, Daniel Szames, Siobhan O’Brien, Stephane Angers, Shana O. Kelley

## Abstract

A major impediment to the characterization of mtDNA repair mechanisms, in comparison to nuclear DNA repair mechanisms, is the difficulty of specifically addressing mitochondrial damage. Using a mitochondria-penetrating peptide, we can deliver DNA-damaging agents directly to mitochondria, bypassing the nuclear compartment. Here, we describe the use of a mtDNA-damaging agent in tandem with CRISPR/Cas9 screening for the genome-wide discovery of factors essential for mtDNA damage response. Using mitochondria-targeted doxorubicin (mtDox) we generate mtDNA double-strand breaks (mtDSBs) specifically in this organelle. Combined with an untargeted Dox screen, we identify genes with significantly greater essentiality during mitochondrial versus nuclear DNA damage. We characterize the essentially of our top hit - WRNIP1 - observed here for the first time to respond to mtDNA damage. We further investigate the mitochondrial role of WRNIP1 in innate immune signaling and nuclear genome maintenance, outlining a model that experimentally supports mitochondrial turnover in response to mtDSBs.

## Introduction

Mitochondrial DNA (mtDNA) bears similarity to its ancestral bacterial genome, as it is circular, polycistronic, and encodes replication and translation machinery including 2 rRNAs and a full complement of 22 tRNAs that can be traced to the endosymbiotic origin of mitochondria^1^. Evolutionary pressure has eliminated redundant processes between nuclear and mitochondrial DNA, leaving mtDNA encoding only 13 OXPHOS proteins, mainly subunits of the electron transport chain (ETC), and forcing mitochondria to rely on nuclear-encoded genes for all other processes including replication, repair, signalling, and maintenance^2,3^. Despite the streamlined structure of this genome, a properly functioning mitochondrial network is so essential that damage to mtDNA can be indirectly sensed as a decrease in respiratory capacity, with the immediate response being an increase in mitobiogenesis^4^.

The types of active DNA repair that ensure mitochondrial genome integrity are still debated^5^. Mitochondria are a hub for oxidative stress, so it follows that base-excision repair (BER) has been best characterized^6^. Conflicting evidence for mitochondrial nucleotide-excision repair (NER) and mismatch repair (MMR) exists in the literature^7,8^. Whether mitochondria participate in double-strand break (DSB) repair remains unclear, with prior work indicating that mtDNA degradation following damage may be favoured instead of direct repair of mitochondrial DSBs (mtDSBs)^9,10^. It has been suggested by Wu *et al*. (2021) that evolutionary pressure has favoured the mitochondrion to produce more mtDNA molecules than necessary for efficient OXPHOS, and to lack mtDNA repair mechanisms so that it may act as a sentinel for cellular genotoxic stress^11^. The high copy number, lack of repair, and increased susceptibility for damage of mtDNA combine to suggest an expendable nature for individual mtDNA molecules. Here, the main goal of our study is to use peptide-based chemical probes developed in our lab to elucidate the cellular response to mtDSBs.

Many mechanisms underlying mtDNA-specific damage response have remained obscure given the difficulty of specifically generating damage in this organelle. Probes used to study nuclear DNA damage are either incapable of accessing the mitochondrial matrix, or access both the mitochondria and the nucleus, leaving questions concerning which responses are mitochondria-specific. Our lab has developed mitochondria-penetrating peptides (MPPs) with cationic and lipophilic character capable of not only crossing the mitochondrial membranes, but also carrying a variety of cargo into the mitochondrial matrix^12,13^. Doxorubicin (Dox) is a chemotherapeutic drug known to cause DSBs in the nucleus. Dox causes poisoning of the topoisomerase II (TOP2) complex after DNA strand breaks have been purposefully used to relieve strand tension^14^. The TOP2-Dox-DNA complex is unable to reconnect the strand break, leaving DSBs throughout the genome. Our previous work demonstrated that Dox can be directed to target mtDNA, causing highly specific mtDSBs^15,16^. As for its potential as a chemotherapeutic, mtDox has been observed to be less cardiotoxic than Dox, and has the potential to evade Dox resistance in cancer cells^17^. Full characterization of the probe (mtDox) is provided in Table 1. This probe has been shown to exclusively damage mitochondrial DNA, while the parent compound primarily damages nuclear DNA.

Previously, we have reported the capacity for mitochondria-targeted agents to probe specific biological processes by using MPP-conjugated DNA-damaging agents in tandem with a limited siRNA screen. This screen targeted 239 known nuclear DNA maintenance factors to identify novel pathways of mtDNA damage response^18^. The siRNA screen, along with a follow up study^19^, was the first to reveal the mitochondrial role of Polθ as an error-prone mtDNA polymerase essential for recovery from oxidative mtDNA damage.

Building on this proof-of-concept screen, we have expanded to a genome-wide CRISPR/Cas9 screen using the TKOv3 library containing 70,948 gRNAs targeting 18,053 genes^20^. While the siRNA screen focused exclusively on known DNA-maintenance genes, this unbiased approach has the power to uncover pathways acting not only directly on mtDNA, but also on DNA damage signalling and cellular rewiring. To validate the compartmentalization of damage, the mtDox screen was conducted in parallel with untargeted Dox, also revealing genes with greater comparative essentially in mtDSB response. Through these sets of chemogenomic screens, including the first to exclusively target mtDNA, we observed the previously unreported requirement of WRNIP1 for the recovery of cells from mtDNA-damage. We further reveal its role as an axis of mitochondrial innate immune response, and its importance for nuclear genome maintenance towards the recovery of mitochondria.

## Results

### CRISPR screen reveals genes essential for mtDSB response

To identify genes with greater essentiality following treatment with mtDox we performed chemogenomic CRISPR/Cas9 screens to compare dropout of gRNAs in a vehicle treated population (DMSO) with cells treated with mtDox or Dox, with the latter compound serving as a control that primarily affects the nucleus (Figure 1a and 1b). The natural fluorescence of Dox was used to visualize its compartmentalized localization (Figure S1a). In negative selection CRISPR/Cas9 screens, gRNAs targeting genes essential for normal proliferation under native conditions will fall out of all populations, however gRNAs that are essential under the damage context will have a more significant dropout in that context, providing evidence that these genes are essential for damage recovery as their KO sensitizes cells to that condition.

**Figure 1:**
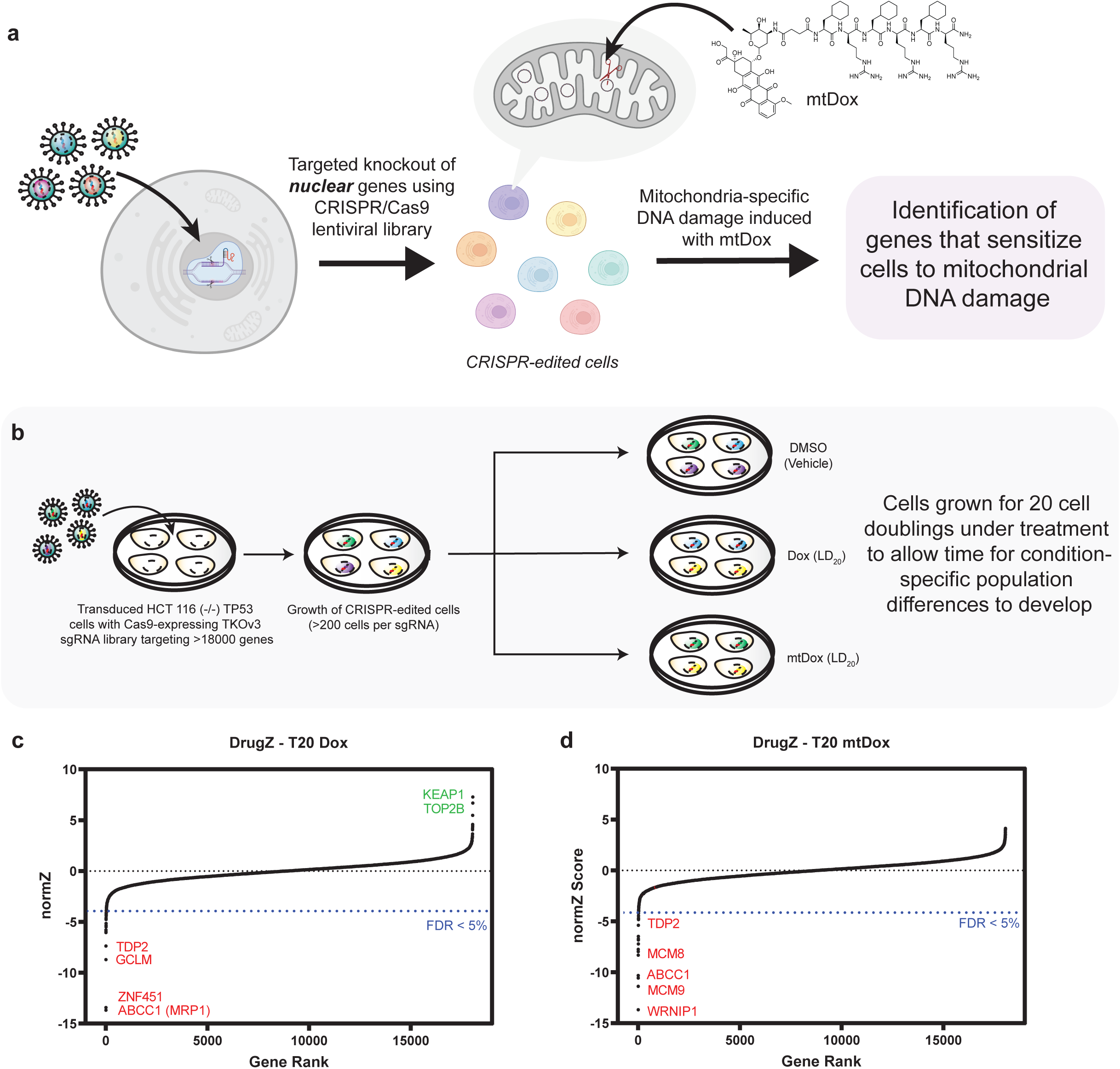
Revealing global response factors to mitochondrial damage using a targeted peptide and a genome-wide CRISPR/Cas9 screen. **(a)** Schematic representation of how biological processes can be revealed by the synergistic application of nuclear CRISPR/Cas9 knockout paired with specific mitochondrial damage using mtDox. **(b)** Overview of the CRISPR/Cas9 screening process with the incorporation of drugs of interest using a negative selection screen over 20 cell doublings. **(c)** DrugZ analysis of untargeted Dox (n = 3 discrete screen replicates). NormZ represented the normalized Z-scored assigned by the algorithm to quantify significance of fold-change from the DMSO vehicle control. A gene with a more negative normZ score indicates a higher susceptibility to death in cells carrying that gene knockout. A more positive normZ score is assigned to gene knockouts which confer resistance to the drug. **(d)** DrugZ analysis of mitochondria-targeted mtDox (n = 3 discrete screen replicates).

For all the screens, HCT116 *TP53*(-/-) cells were treated with a low level of compound designed to kill no more than 20% of cells relative to DMSO control (LD_20_) (Figure S1b), as treating with higher concentrations can lead to off-target effects related to cell death. HCT116 cells were chosen based on previous evidence showing their sensitivity to mitochondrial disruption in CRISPR/Cas9 essentiality screens^21^. To pressure the cells to rely on mitochondrial OXPHOS over glycolysis for energy, a reduced glucose concentration was used in the media and supplemented with pyruvate. Some DNA-damage related genes may be masked if p53 is present therefore, to bypass this damage-control mechanism and increase the efficiency of CRISPR/Cas9 knockout, we used a cell line with p53 removed^22,23^.

Following TKOv3 lentiviral library infection targeting 18,053 genes, and a brief selection period, the cells were passaged and treated for 20 cell doublings to allow significant genetic drift of the populations. Library gRNA sequences were excised at the beginning and end of the screen and sent for deep sequencing to measure the abundance of individual gRNAs in each population. The MAGeCK count algorithm was used to map reads to gRNAs^24^, and the drugZ algorithm^25^ was used to assign normalized Z-scores (normZ) to differential fold-changes of gRNAs between populations (Figure 1c and 1d). As a control, the screens were also monitored for expected dropout of previously categorized essential genes using the BAGEL algorithm (Figure S2a)^21^.

The analysis of the sequencing data from the screen indicated that our approach was able to reveal context-dependent essential genes, as the top gene hits for Dox were *ABCC1*, *ZNF451*, *GCLM*, and *TDP2* (Figure 1c). *ABCC1* encodes for multidrug resistance-associated protein 1 (MRP1), a protein whose over-expression is linked to doxorubicin resistance in chemotherapy^26^. *ZNF451* and *TDP2* encode proteins involved in the disassembly of the stalled topoisomerase II complex^27^. Mice lacking *GCLM* have been shown to overexpress carbonyl reductase 3 (CBR3) leading to increased metabolism of doxorubicin to its more cardiotoxic form of doxorubicinol^28^.

On the other end of the spectrum, *TOP2B* knockout was associated with increased survival in Dox-treated cells. This observed resistance to Dox in *TOP2B* knockout cells falls in line with evidence that if the TOP2B protein is not present, Dox is unable to poison the TOP2 complex and generate DSBs^29^. Another Dox resistance gene observed was *KEAP1.* The KEAP1 redox sensor protein negatively regulates NRF2 (nuclear factor erythroid 2-related factor 2) which activates antioxidant response elements (ARE)^30,31^. Exposure to Dox has been observed to cause autophagic degradation of KEAP1 in response to Dox-generated ROS. Deletion of KEAP1 would lead to the constitutive stabilization of NRF2 and expression of AREs. Stable over-expression of NRF2 has been observed to cause resistance to chemotherapeutic agents such as cisplatin, doxorubicin (Dox), and etoposide^32^. Our Dox screen results are similar to a screen published by Olivieri *et al*., which also observed *ABCC1*, *ZNF451*, and *TDP2* as top essential genes, and *TOP2B* and *KEAP1* as top resistance knockout genes^23^, validating the design and execution of our approach.

The top differentially essential genes in the mtDox-treated population include *WRNIP1*, the *MCM8/9* and *C17orf53* (*HROB*) complex, as well as *RECQL5* (Figure 1c). *ABCC1*, *ZNF541*, and *TDP2* were also essential in the mtDox-treated population, however comparison of the top hits between populations (*FDR* < 0.05) revealed unique sets of hits to each treatment condition (Figure 2a and 2b). GO Panther analysis^33^ was used to analyze the essential genes from each population with a *p*-value assigned by drugZ as < 0.05 (Figure 2c). While mitochondria-related processes do not initially stand out in the mtDox-treated population, comparing the differential top hits between mtDox and Dox highlights an enrichment of distinctly mitochondrial genes such as mitochondrial translation and mitochondrial gene expression and complex assembly.

**Figure 2:**
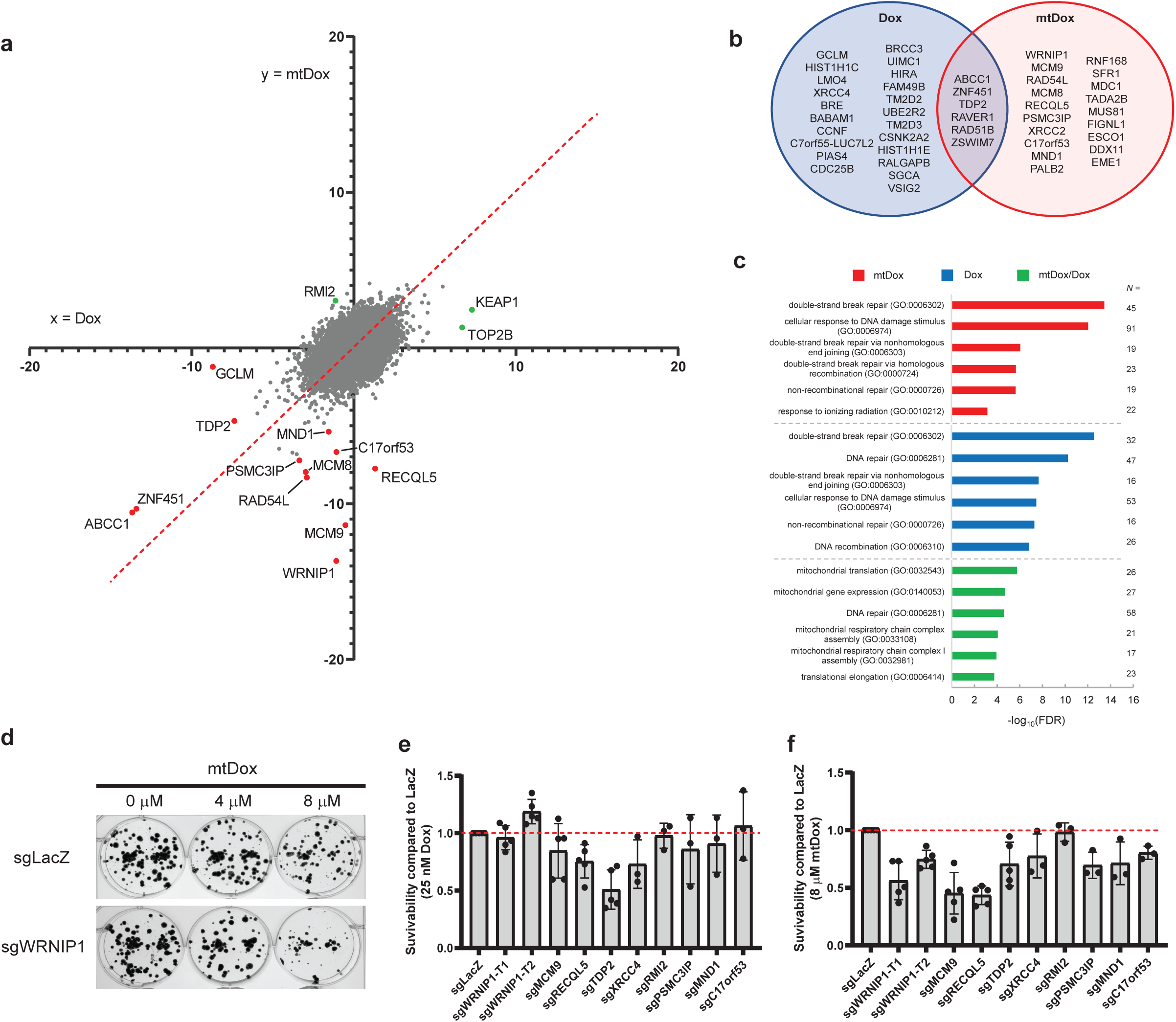
Comparative results, data trends and validation between Dox and mtDox screens. **(a)** Direct comparison of normZ scores between Dox (x-axis) and mtDox (y-axis). Red line drawn through x=y indicates genes with equal essentiality in both Dox and mtDox treated conditions. Points below the line represent genes which are more essential during mtDox damage compared to Dox. **(b)** Venn diagram representing the genes which had an *FDR* < 0.05 attached to their normZ value for each drug. **(c)** Biological process analysis of genes with *p*-values < 0.05. **(d)** Pictorial representation of clonogenic analysis for susceptibility of WRNIP1 knockout cells to mtDox treatment. **(e)** Quantification of clonogenic analysis using 8 μM mtDox for select mtDox screen hits (for e and f, n = 5 for sgWRNIP1 through sgTDP2, n = 3 for sgXRCC4 through sgC17orf53). RMI2 represents a control gene whose knockout did not lead to increased susceptibility to mtDox based on the CRISPR/Cas9 screen. Data represented as mean ± s.d.

The CRISPR/Cas9 screen was validated by testing the sensitivity of cells harboring knockout of the top hits identified in the screen to Dox or mtDox using a clonogenic assay^23^ (Figure 2d and S2b). Ten targets were chosen for validation including two distinct sgRNA sites for *WRNIP1,* annotated as T1 and T2 for Target 1 and 2. *XRCC4* and *TDP2* were selected as control genes observed to be essential in both Dox and mtDox-treated populations. *XRCC4* and *TDP2* have previously been characterized for their roles in mitochondria using mtDox^18,34^. *RMI2* was observed to be non-essential in the presence of mtDox according to the screen and was therefore used as a negative control for mtDox clonogenic validation. Overall, the clonogenic assays recapitulate the data captured in the CRISPR/Cas9 screens.

A more comprehensive clonogenic analysis is provided (Figure S3) including additional Dox and mtDox concentrations, a comparison with wild-type (WT) HCT116 *TP53*(+/+) cells, the use of high glucose concentration on HCT116 TP53(-/-) cells, and the use of a different cell line, RPE1 *TP53*(-/-). Sensitivity to mtDox in WRNIP1 knockout cells was observed in all conditions tested except for high glucose conditions in which cells may rely on non-mitochondrial sources of energy.

### The role of WRNIP1 in mitochondrial damage response

WRNIP1 (WRN interacting protein 1), is an AAA+ ATPase associated with a variety of DNA-repair mechanisms^35^. Most of the observed functions of WRNIP1 involve its localization to DNA and recruitment of repair factors. WRNIP1 recruits FANCD2 to sites of interstrand crosslinks and interacts with RAD51 and BRCA2 to promote stabilization of stalled replication forks^36^. WRNIP1 plays a role in genome maintenance through interaction with PCNA and POLD1 as well as through controlling the levels of PRIMPOL in the cell^37^. WRNIP1 has also been associated with innate mitochondrial antiviral signalling (MAVS) through stabilization of the interaction between RIG-I and cytoplasmic dsRNA, an essential component of immune activation through this pathway^38^.

Notably, while WRNIP1 was a strong hit in the mtDox screen, its knockout did not significantly affect the survival of cells treated with Dox or other DNA damaging agents. Compared to screens conducted by Olivieri *et al*., WRNIP1 does not appear as a hit for any of the 27 genotoxic agents tested including DNA-damaging agents: cisplatin, H_2_O_2_, UV, IR, methyl methanesulfonate (MMS), bleomycin, and etoposide^23^. This evidence highlights the advantage of using mtDox to study compartmentalized mitochondrial damage for WRNIP1 involvement in mtDNA damage response, as its essentiality is only observed under mitochondria-specific damage. Additionally, we have conducted CRISPR/Cas9 screens using other MPP-conjugated compounds such as mtOx, a thiazole orange-based oxidizing agent (Figure S4a). Comparing the hits with a *p*-value of < 0.05 between the mtOx (*N* = 842) and mtDox (*N* = 825) screens, there are only 37 gene hits which appear in both screens, indicating that the peptide is likely not a significant source of off-target gene hits. WRNIP1 essentiality was also not observed in the mtOx screen.

We compiled a list of observed cellular consequences to mtDox treatment to help elucidate the potential role of genes necessary for recovery from mtDox damage (Table 1, Figure S4). In general, we hypothesize a model that supports mitochondrial turnover as opposed to mtDSB repair, as we do not observe an increase in mtDNA mutations from long-term mtDox-treatment but do observe an increase in mitochondrial mass as a potential compensatory mechanism.

To investigate why WRNIP1 is essential for cell survival following mtDox treatment, monoclonal cell lines were generated using CRISPR/Cas9 which contained no observable WRNIP1 protein (Figure 3a and S5a). The cell lines can be grown stably for long periods of time, indicating that WRNIP1 is not essential under normal conditions. To assess whether WRNIP1 interacts with mtDNA directly, we performed a proteinase K assay to characterize WRNIP1 localization (Figure 3b and S5b). For this assay, mitochondria were first stringently isolated using sucrose-gradient ultracentrifugation. Next, isolated mitochondria were either subjected to no treatment, proteinase K, or proteinase K with SDS. Proteins within the mitochondria are protected from degradation by proteinase K for the duration of the experiment. WRNIP1 does not appear to localize to either the mitochondrial matrix like TFAM nor the outer mitochondrial membrane like BCL-XL, therefore WRNIP1 is not participating in direct stabilization or repair of mtDNA since it does not access the mitochondrial matrix.

**Figure 3:**
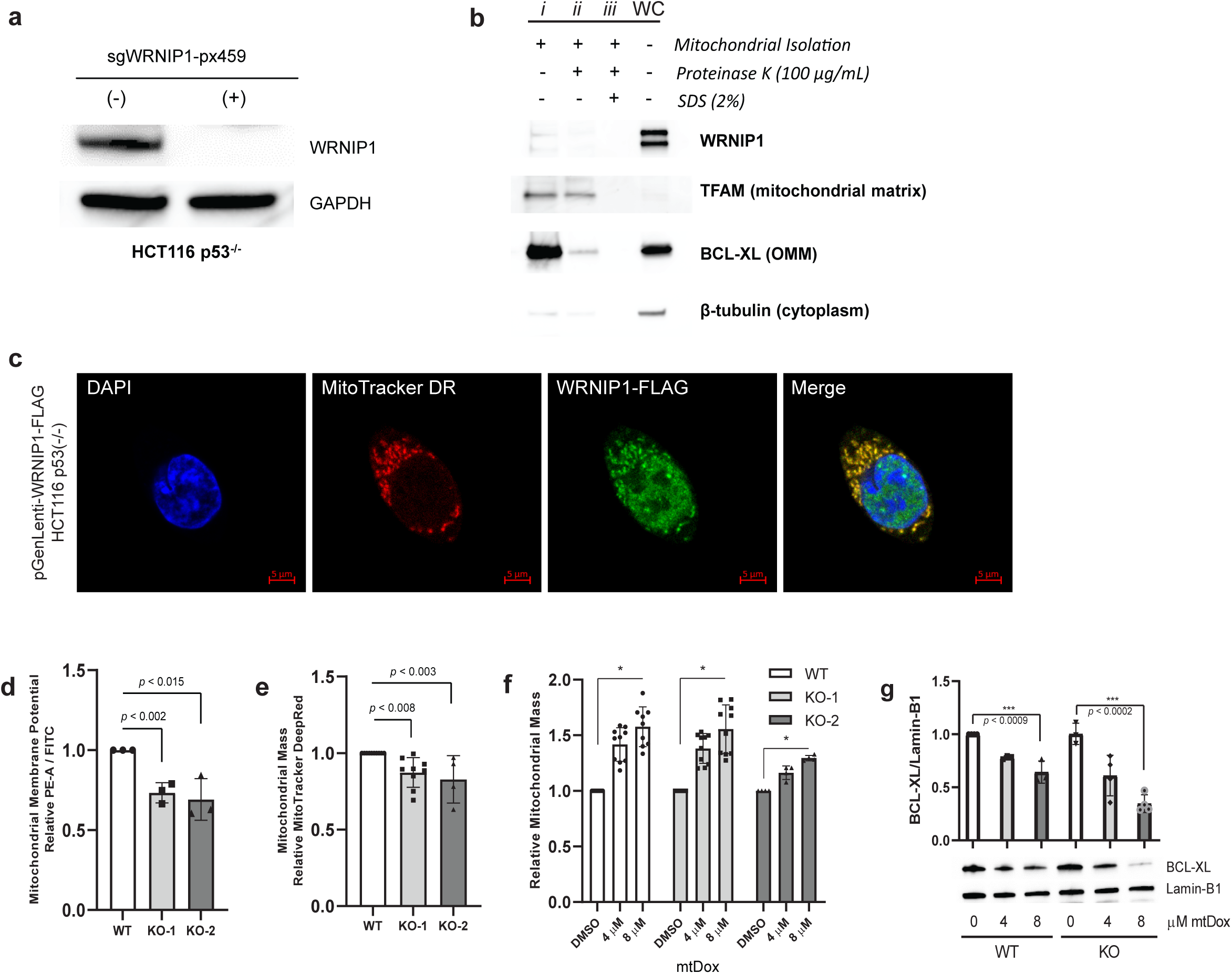
Characterization of the mitochondrial role of WRNIP1. **(a)** Monoclonal WRNIP1 knockout (KO) cell line using transient px459 plasmid-based CRISPR/Cas9 knockout. **(b)** Proteinase K protection assay for mitochondrial localization. WC = whole cell, *i* = purified sucrose-gradient ultracentrifugation of mitochondria, *ii* = *i* + 100 μg/mL Proteinase K for 30 mins, *iii* = *ii* + 2 % SDS. WRNIP1 is not enriched in the mitochondrial fraction like TFAM and does not appear inside the mitochondria. **(c)** Confocal microscopy localization of WRNIP1-FLAG in fixed cells stained with MitoTracker Deep Red (633 nm) and DAPI (nucleus). **(d)** Flow cytometry analysis of basal mitochondrial membrane potential using JC-1 dye between WT cells and two monoclonal WRNIP1 KO lines. Data represented as aggregate polymer (PE-A) over monomer (FITC) to normalize for mitochondrial mass (n = 3). **(e)** Flow cytometry analysis of mean basal mitochondrial mass compared between WT and two distinct monoclonal WRNIP1 KO lines using the APC channel (n = 9 for WT and KO-1, n = 4 for KO-2). **(f)** Flow cytometry analysis of mean mitochondrial mass using MitoTracker Deep Red. Cells were treated with mtDox for 24 hours before staining and immediate live analysis using the APC laser channel (n = 9 for WT and KO-1, n = 4 for KO-2, \**p* < 0.0001). **(g)** Western blot measuring anti-apoptosis protein BCL-XL with increasing concentrations of mtDox in parent (WT) and WRNIP1 knockout (KO) cell lines over 72 hours (n = 3). All *p*-values on graphs determined using unpaired *t*-test. Data represented as mean ± s.d.

For WRNIP1 localization, we performed confocal microscopy with a generated WRNIP1-FLAG expressing cell line using a pGenLenti construct with a CMV promoter. Lentiviral infection followed by puromycin selection was used to incorporate expression (Figure S5a). Using this over-expression cell line and an anti-FLAG antibody, strong mitochondrial association was observed for WRNIP1-FLAG potentially due to its reported association with MAVS, found on the OMM (Figure 3c)^38^. Using an overexpression vector increased our chances of picking up the general localization of WRNIP1 using microscopy. This interaction was not observed using the proteinase K assay alone, also potentially due to the transient nature of the interaction between WRNIP1 and MAVS, and the lack of direct interaction between WRNIP1 and the OMM.

Indicators of mitochondrial health are mitochondrial polarization and mitochondrial mass. We used flow cytometry to measure the potential of the mitochondrial membrane using JC-1 dye and mitochondrial mass using MitoTracker Deep Red. Interestingly, cells lacking WRNIP1 appear to have lower mean basal membrane potential, and lower mean mitochondrial mass (Figure 3d and 3e). Due to its inherent fluorescence crossover with JC-1 dye, we were unable to measure the effect of mtDox on membrane potential but could use MitoTracker Deep Red to monitor mitochondrial mass. We observed a significant and concentration-dependent increase in mean mitochondrial mass with the introduction of mtDox (Figure 3f). The knockout of WRNIP1 did not consistently alter the observation of increased mitochondrial mass in the two monoclonal KO lines tested.

We performed a series of tests using the WRNIP1 knockout cell line to assess mitochondria-mediated cell death. BCL-XL is a mitochondria-associated anti-apoptotic protein, and the presence of BCL-XL prevents release of mitochondrial contents into the cytoplasm and triggering of caspase-mediated cell death^39^. We monitored BCL-XL expression after 72 hours of 4 and 8 μM mtDox treatment (Figure 3g). The anti-apoptotic protein is reduced in a mtDox-dependent manner in both the WT and WRNIP1 KO cell lines, however cells lacking WRNIP1 had more significant reduction in BCL-XL expression, indicating increased mitochondria-mediated apoptosis with WRNIP1 KO. We probed for expression of LAMP1, LC-3, and OPA1 to assess the effect of mtDox and WRNIP1 on autophagy and mitochondrial network dynamics but did not observe significant differences in their expression (Figure S5c). WRNIP1 KO does not affect basal levels of mtDNA, nor the extent of mtDNA copy number reduction by mtDox treatment (Figure S5d and S5e). Probing for WRNIP1 expression over time, we did not observe any changes associated with mtDox treatment (Figure S5f). We performed a Seahorse mitochondrial stress test and observed decreased oxygen consumption rate (OCR) in WRNIP1 KO cells at basal conditions as well as ATP production conditions. Treatment with 8 μM mtDox for 6 hours in both WT and WRNIP1 KO cells caused a reduction in OCR levels for basal, proton leak, ATP production, and maximal respiration conditions (Figure S6).

Overall, WRNIP1 knockout appears to cause a reduction in mitochondrial membrane potential, mitochondrial mass, and mitochondrial respiration, however the mechanism of mtDox induced death remains unclear.

### mtDox-activated innate immune response requires WRNIP1

The release of mitochondrial contents into the cytoplasm elicits an immune response thought to be related to their evolutionary origin as bacteria, with their contents mimicking those of a pathogenic invader^40^. The mitochondria-derived damage-associated molecular patterns (DAMPs) include: ATP, succinate, N-formyl peptides, cardiolipins, TFAM and mtDNA^41^. Free cytoplasmic mtDNA is associated with aging, chronic inflammatory diseases, cardiac ischemia, and a growing number of diseases^42^.

The cytoplasmic sensors for mtDNA are generally thought to be cGAS/STING and TLR9, however a recent study using mitochondria-targeted TALENs to cause mtDSBs concluded that the main sensor triggered by this type of damage was the RIG-I/MAVS pathway^43^. While the primary purpose of RIG-I is the recognition of viral dsRNA, an increase of mitochondrial dsRNA released into the cytoplasm was observed following the generation of mtDSBs that was sufficient to activate the RIG-I mediated innate immune response.

Prior work using RNAi and yeast two-hybrid screening identified WRNIP1 as a key player in the activation of MAVS through directing RIG-I to the TRIM14/MAVS docking platform^38^. The activation of MAVS was dependent on the presence of WRNIP1 during VSV infection. Another study used the mtDox compound to observe its effect on immune response^44^. mtDox alone was able to activate a variety of genes associated with immunity as evidenced by their increased RNA-level expression. In addition, mtDox increases priming of treated cells for immunogenic cell death as observed by monitoring cell-surface levels of calretriculin^17^. With these key studies in mind, we hypothesized that the presence of WRNIP1 as the top hit in the mtDox CRISPR screen is partly due to its essentiality in activating the innate immune response leading to nuclear sensing of mitochondrial damage and replacement with viable mitochondria.

A schematic of the proposed innate immune pathway is provided (Figure 4a). Treatment with mtDox causes a release of DAMPs into the cytoplasm including dsRNA and which activates the RIG-I/MAVS innate immune pathway through WRNIP1. This immune activation signals a mitochondrial health checkpoint response in the nuclear compartment. To connect the missing pieces of this pathway, we aimed to demonstrate that mtDox alone could trigger association between WRNIP1 and RIG-I and that the downstream immune activation by mtDox is dependent on the presence of WRNIP1.

**Figure 4:**
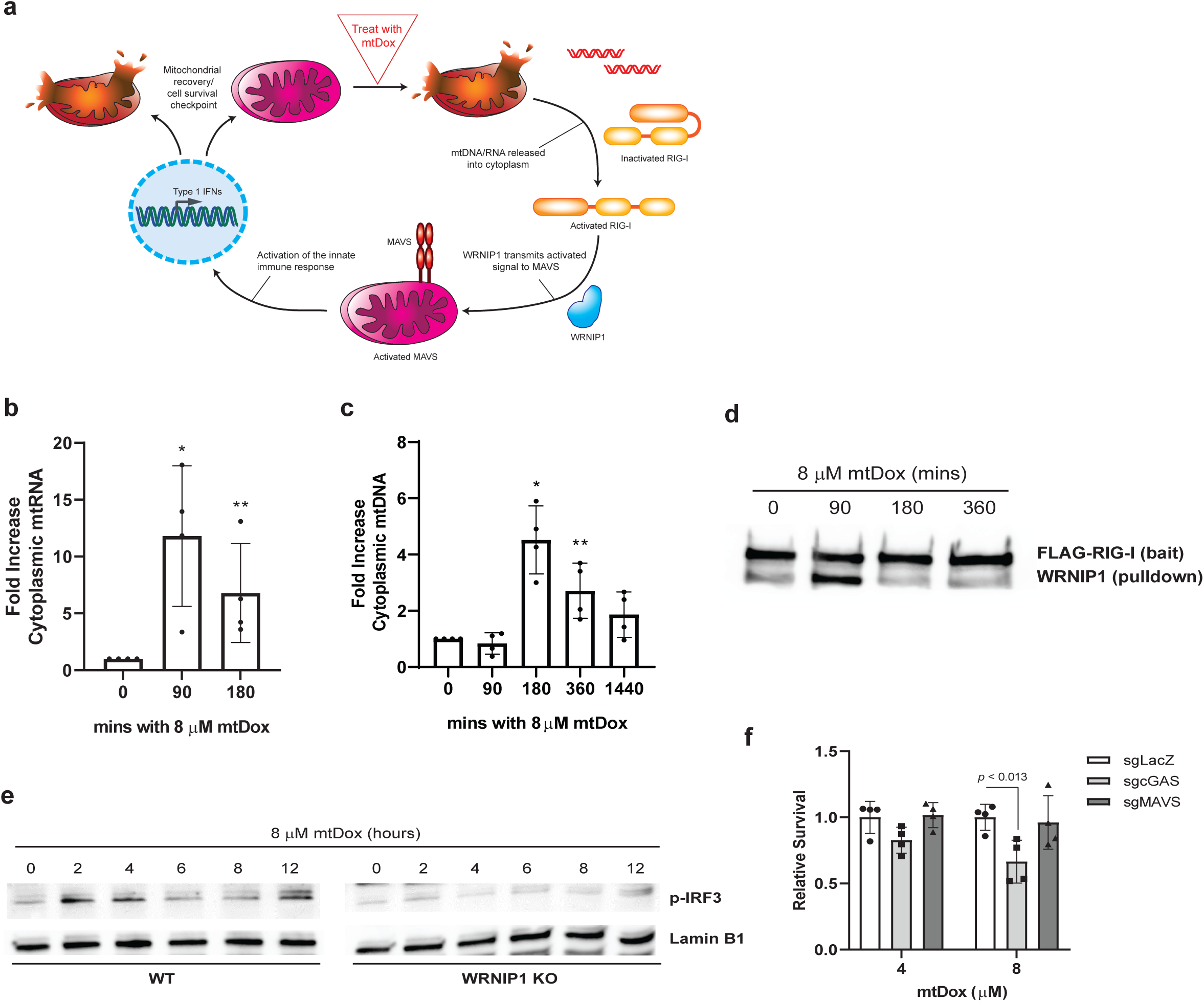
Mapping the function of mtDox and WRNIP1 in innate immune activation. **(a)** Schematic representation of the proposed cellular response to mtDox damage. mtDNA damage caused by mtDox leads to mitochondrial herniation followed by release of mitochondrial contents into the cytoplasm, sensed through RIG-I innate immune pathway. Functional WRNIP1 transports activated RIG-I to MAVS which stimulates immune response activation, leading to nuclear recognition of damage. **(b)** Digitonin based extraction followed by qPCR measurement of mtDNA released into the cytoplasm over time following 8 μM mtDox treatment (n = 4, \**p* < 0.0011, \*\**p* < 0.013). **(c)** Digitonin based extraction followed by RT-qPCR measurement of mtRNA released into the cytoplasm over time (n = 4, \**p* < 0.013, \*\**p* < 0.037). **(d)** Western blot of RIG-I-FLAG co-immunoprecipitation displaying time-course interaction of RIG-I with WRNIP1 following mtDox treatment. Anti-FLAG M2 magnetic beads were used to pull down RIG-I-FLAG transfected lysates, and blot was counter-stained with antibodies for FLAG and WRNIP1 (n = 3). **(e)** Time-course Western blot for phosphorylated IRF3 (p-IRF3) indicating downstream innate immune activation. Performed with parent (WT) and WRNIP1 knockout (KO) cells lines (n = 3). **(f)** Clonogenic analysis of relative survival following mtDox treatment between LacZ, cGAS, and MAVS knockout (n = 4). All *p*-values determined using unpaired *t*-test. Data represented as mean ± s.d.

As the first step in the proposed pathway, we investigated whether mtDox would elicit a release of mtRNA into the cytoplasm which could act as a trigger for MAVS activation. We used a digitonin-based cytoplasmic extraction method and RT-qPCR for mtRNA analysis. We observed a significant increase in mtRNA in the cytoplasmic fraction peaking at 90 minutes post-treatment with 8 μM mtDox (Figure 4b). We also tested whether mtDNA was released into the cytoplasm and observed its presence following treatment with 8 μM mtDox using qPCR with mtDNA specific probes (Figure 4c).

To demonstrate that WRNIP1 associates with RIG-I in the presence of mtDox, we set up a time-course co-immunoprecipitation (CoIP) using a RIG-I-FLAG construct as bait. The cells expressing RIG-I-FLAG were treated for 0, 90, 180, and 360 minutes with mtDox. The whole-cell lysates were pulled down by anti-FLAG magnetic beads. An interaction between WRNIP1 and RIG-I was observed up at 90 minutes following mtDox treatment (Figure 4d) indicating that the interaction is time-sensitive and mtDox dependent.

To further probe this pathway, we tested whether mtDox could activate the innate immune response. Downstream activation of MAVS can be tracked by monitoring the phosphorylation of IRF3 over time using VSV infection^38^. We adapted this approach with time points of 0-, 2-, 4-, 6-, 8-, and 12-hours post-treatment with mtDox (Figure 4e). The experiment was conducted in parent (WT) and WRNIP1 knockout cell lines (KO). The addition of mtDox caused an increase in IRF3 phosphorylation between hours 2 and 4 in WT cells. This increase in IRF3 phosphorylation was attenuated in the WRNIP1 KO cells, indicating that WRNIP1 is required for the cells to respond to mtDox through this immune pathway, supporting its role as essential for mitochondrial damage response.

To investigate whether this innate immune pathway is the avenue for WRNIP1 essentiality following mtDox treatment, a clonogenic assay was set up to compare survivability between sgLacZ, sgcGAS, and sgMAVS. Interestingly, while we did not observe sensitivity to mtDox treatment in MAVS KO cells, we did observe a significant decrease in survival from cGAS KO cells under mtDox treatment (Figure 4f). Having observed the release of mtRNA and mtDNA following mtDox treatment, it is possible that both the MAVS and cGAS pathways are activated by mtDox-induced damage. As WRNIP1 has only currently been observed to interact with the MAVS pathway of innate immune activation, and MAVS KO does not lead to mtDox sensitivity, the mechanism for WRNIP1-induced essentiality for recovery from mtDox treatment is likely not driven by its participation in the innate immune response. Future studies could however be used to investigate whether WRNIP1 also acts within the cGAS/STING pathway.

### WRNIP1 knockout leads to genomic instability

To explore the role of WRNIP1 as a nuclear genome maintenance factor, we tested whether the long-term knockout of WRNIP1 leads to increased basal levels of nuclear DNA damage^36,37^. Loss of a maintenance factor such as WRNIP1 can lead to replication stress and the accumulation of unrepaired DNA. We used the comet assay to compare untreated parent (WT) versus WRNIP1 knockout (KO) cells for evidence of nuclear DNA damage. In this assay, single cells trapped in agarose are subjected to an electric current, which draws out damaged particles of DNA into a comet-tail trailing the nucleus. We observed the comet tail motif in KO cells but did not detect significant basal DNA damage in WT cells (Figure 5a).

**Figure 5:**
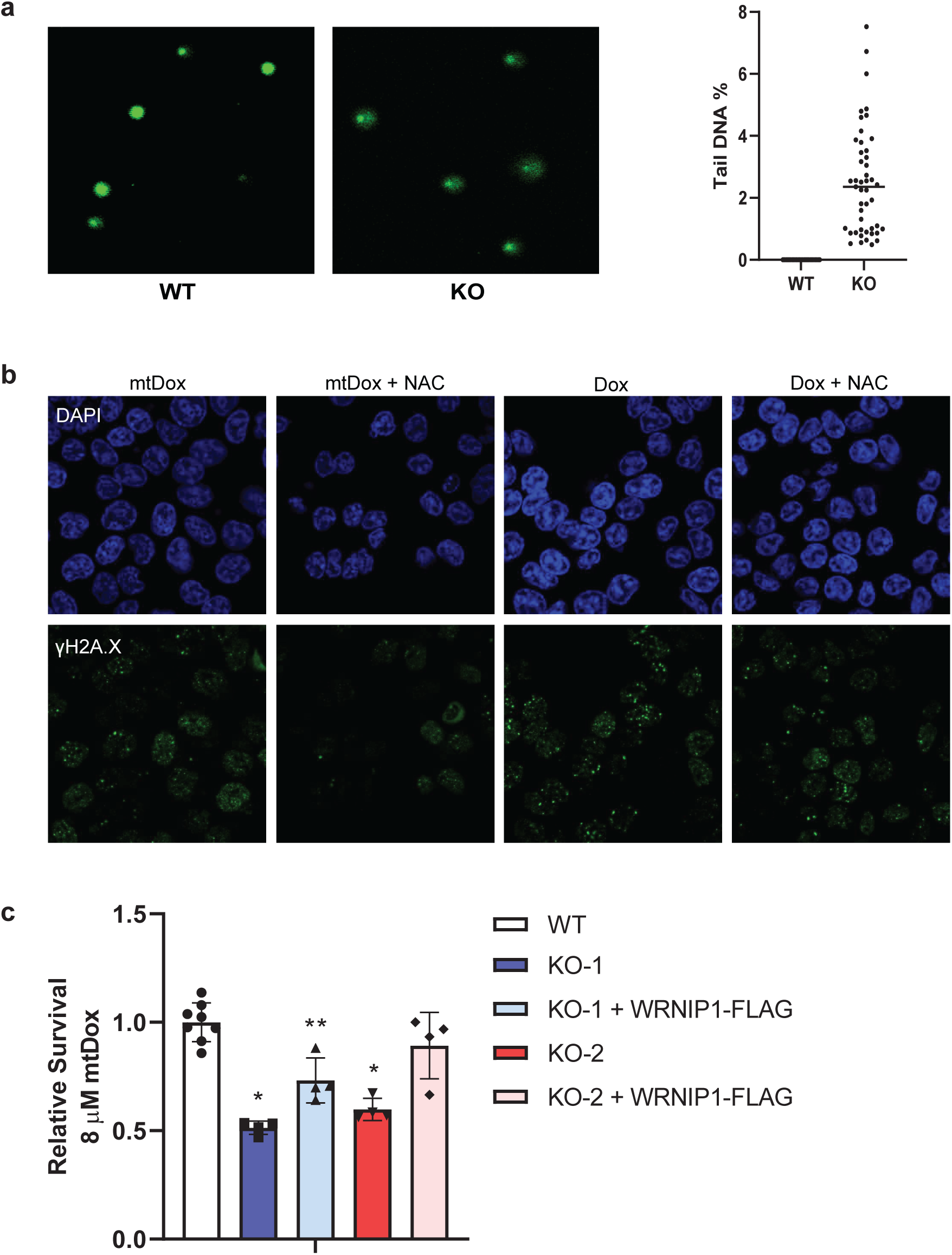
Loss of WRNIP1 leads to genomic instability. **(a)** Comet assay detecting basal nuclear DNA damage in WRNIP1 KO cells versus WT. Quantification of comet tail DNA % is measured by ImageJ using microscopy images (n = 3 discrete replicates). Data points represent each individual cell analyzed. **(b)** Detection of γH2AX foci representing nuclear DNA damage in Dox and mtDox treated cells. Microscopy images comparing 24-hour treatment of HCT116 *TP53*(-/-) cells with 8 µM mtDox and 25 nM Dox with and without 10 mM *N*-acetyl cysteine (NAC) antioxidant treatment. Moderate γH2AX detection in mtDox treated cells is attenuated by antioxidant treatment, while cells treated with Dox still display pronounced nuclear DNA damage in the presence of antioxidant. **(c)** Clonogenic survival comparing representative survival of cell lines treated with 8 μM mtDox. KO-1 and KO-2 represent two monoclonal WRNIP1 KO lines of HCT116 *TP53*(-/-) (WT). The reintroduction of a WRNIP1 construct shows partial but incomplete recovery of the monoclonal KO lines to WT standard (n = 4, *p*-values are compared to WT, \**p* < 0.00001, \*\**p* < 0.009) All *p*-values determined using unpaired *t*-test. Data represented as mean ± s.d.

We aimed to investigate whether mtDox was contributing to increased genomic instability through an indirect mechanism acting on the nucleus. While mtDox itself does not access nuclear DNA, it does generate significant amounts of cellular reactive oxygen species (ROS) (Figure S4c). Observing γH2A.X puncta in the nucleus, we compared nuclear DNA damage caused by Dox versus mtDox (Figure 5b). Nuclear DNA damage is observed following 24 hours mtDox and Dox treatment – however to a lesser extent through mtDox treatment. To demonstrate that this damage is the indirect result of ROS produced by mtDox in the nucleus, we co-treated the cells with a potent antioxidant *N*-acetyl cysteine (NAC) and observed ablation of the mtDox-induced nuclear damage but not the direct Dox-induced damage.

We propose that the increase in basal nuclear DNA damage caused by WRNIP1 KO, combined with accumulated nuclear ROS damage, leads to a reduction in the ability of the cell to replace mitochondrial proteins and mitochondrial maintenance factors encoded in the nucleus. The combination of nuclear DNA stress caused by the loss of WRNIP1 with targeted damage of mitochondria by mtDox provided the unique conditions to draw WRNIP1 as the top hit in our CRISPR/Cas9 screen.

Another point of evidence for the accumulation of nuclear damage in WRNIP1 KO cells leading to increased sensitivity to mtDox is the inability for this phenotype to be rescued by the addition of a WRNIP1 construct (Figure 5c). While there is moderate rescue of the monoclones, the rescue is incomplete, particularly in KO-1 which is a monoclonal line that has been cultured for approximately double the number of passages as KO-2. A final point of evidence for the time-dependent accumulation of nuclear DNA damage being necessary to observe WRNIP1 essentiality is that sensitivity to mtDox is not observed to the same extent in shorter time course survivability experiments (Figure S7). Lastly, since we did not observe other members of the MAVS pathway in our mtDox screen hits, but did observe many members of nuclear DNA repair, nuclear genome integrity is the likely driver for cell survival following mtDNA damage.

## Discussion

The specificity of the MPP delivery vector to the mitochondrial compartment has made possible this first functional chemogenomic CRISPR/Cas9 screen probing for essential factors following mtDNA-specific damage. Through this chemically designed screening approach we were able to reveal the previously uncharacterized essentiality of WRNIP1 during mtDSB damage response. As WRNIP1 was not observed to localize inside the mitochondria itself and its mechanism of response appears to be indirect from the origin site of mtDNA damage, this essentiality may not have been discovered through classical proteomic screening of mitochondrial interactors.

In summary, WRNIP1 knockout cells were observed to have lower basal mitochondrial membrane potential, mitochondrial mass, and respiration capacity, however its knockout does not seem to affect the extent of direct mtDNA damage caused by mtDox. The essentiality of WRNIP1 for mitochondrial repair appears to be a combination of an innate immune response compounded by its role in nuclear DNA integrity which then negatively impacts the generation of new mitochondria. We have also presented evidence of an mtDox activity model which relies on synergistic mtDNA damage and ROS-acquired nuclear damage leading to cell death.

We aimed to develop a unifying mechanism for WRNIP1 involvement in mitochondrial repair however we instead uncovered a several potential avenues of indirect responses to mtDNA damage through exploring the native response to mtDox treatment, and the conditions in which WRNIP1 is necessary for the expected cellular response. Part of the difficulty in finding direct repair pathways may be due to redundancy in mitochondrial response elements, or as Wu *et al*. (2021) suggests, it may be evolutionarily advantageous for mtDNA damage to run its course, act as a signal for genotoxic damage, and prime the nuclear genome to enhance nDNA repair and surveillance^11^. The cGAS/STING pathway is thought to be integral to the recognition of mtDNA in the cytoplasm, and indeed we have shown that it is necessary for recovery from mtDox damage. We did not however observe the cGAS/STING pathway nor the MAVS pathways as hits in our mtDox whole-genome CRISPR screen which draws attention to a major limitation of this approach for discovery of gene function. Because the drugZ algorithm compares endpoint gene pools between conditions, any genes with essentiality under normal conditions may not present as significantly more essential under the treatment condition. The lack of observed canonical mitochondrial genes as top hits in the mtDox screen may also be attributed to this phenomenon. The use of CRISPR/Cas9 screening for the discovery of mitochondrial repair genes may be inadvertently selecting for accessory genes which are only recruited under specific conditions. This can also be viewed as an advantage however because these genes would not be discovered through more direct methods and the end result is a more holistic view of the processes involved in damage response. Taking all observations into account, including the necessity of cGAS, the activation of ISGs, the prevalence of nuclear repair factors as hits in our mtDox screen (including WRNIP1, MCM8/9, and RECQL5) the overall picture suggests heightened genome surveillance and integrity is necessary following activation of ISGs by mtDox through innate immune mechanisms. The involvement of WRNIP1 in both immune activation and nuclear genome integrity can explain how it became the top hit in our CRISPR screen. The extent to which WRNIP1 essentiality is based on its direct role in innate immune activation versus its absence simply priming the nuclear genome for ROS assault remains to be explored. Further investigation can also be placed in observing whether the participation of WRNIP1 in an active innate immune response limits its availability for nuclear DNA maintenance.

Building on the discussion of context-dependent functionality, we chose a p53-null model of HCT116 for our screens to increase the efficiency of uncovering DNA-damage repair genes however we would be remiss if we did not address the importance of p53 for mitochondria. For nuclear DNA damage, p53 is involved in cell cycle arrest and promotion of apoptosis in non-viable cells and is arguably one of the most important transcription factors in tumour progression^45,46^. As for its mitochondrial role, p53 has been observed to promote mitochondrial genome maintenance and translocates to the mitochondria to promote apoptosis^47,48^. p53 has also recently been associated with MAVS in promoting DNA-damage induced cell death through etoposide treatment^49^. MAVS overexpression induced apoptosis in p53 WT but not p53 KO cells treated with etoposide. While our screen may not recapitulate the environment of a healthy cell response, aberrations in p53 are hallmarks of cancer progression, and our data can serve to reflect these conditions.

While this screen was limited by its use of a fast-growing p53-modified cancer cell model to accommodate the coverage of the CRISPR/Cas9 screen library, this also allows the rapid screening of mitochondria-targeted compounds. The scope provided by this experimental approach opens the door to the examination of a variety of types of mitochondrial damage. Further studies will focus on a catalog of MPP-conjugated damage agents as well as a continuation into the exploration of the expansive hits from this screen to generate a comprehensive directory of mitochondrial damage response.

## Acknowledgments

The project described was supported by Grant Number R01GM116886 from NIGMS. Its contents are solely the responsibility of the authors and do not necessarily represent the official views of the NIH. We acknowledge the support of the Natural Sciences and Engineering Research Council of Canada (NSERC), [funding reference number CGSD3-518428-2018]. We acknowledge The Centre for Pharmaceutical Oncology at the Leslie Dan Faculty of Pharmacy for the use of its resources and equipment. The TKOv3 virus was a gift from the Angers Lab with viral prep performed by S. Lin. NGS of the CRISPR screens was conducted by the Lunenfeld-Tanenbaum Research Institute Sequencing Facility. NGS of the isolated mtDNA was conducted by the Donnelley Sequencing Centre.

## Author Contributions

T.S. performed the experiments and most data analysis with help from P.D. and D.S. CRISPR screen analysis was performed by S.O. who also helped with screen design. CRISPR expertise, library and manuscript editing was provided by S.A. Manuscript was written and edited by T.S. and S.O.K.

## Declaration of Interests

The authors declare no competing interests.

## Tables

**Table 1:** Observed general effects of mtDox on cells.

| Characteristic | Explanation | Evidence |
| --- | --- | --- |
| **mtDox causes cell death within 72 hrs** | Dose-response curves show cell death after 72 hours mtDox treatment. | Figure S1b |
| **mtDox-damaged mtDNA is replaced and not repaired.** | If mtDSBs are repaired, evidence should exist of error-prone (NHEJ) or microhomology-mediated end joining (MMEJ). We sent isolated mtDNA for deep sequencing and did not observe increased mtDNA mutation rate. | Figure S4b |
| **mtDox causes a reduction in mtDNA copy number** | qPCR analysis of mtDNA reveals that mtDNA is being degraded at 6 hours post-treatment. | Figure S4b |
| **mtDox causes an increase in cellular ROS** | Cellular ROS is a significant factor generated by mtDox but not Dox using concentrations employed in clonogenic validation experiments. | Figure S4c |
| **mtDox causes increased mitochondrial mass** | As measured by MitoTracker Deep Red.<br>Mitochondrial mass is increased potentially as a compensatory mechanism for the loss of mtDNA. | Figure 3f |
| **mtDox releases nucleic acids into the cytoplasm** | Damage caused by mtDox causes mitochondrial herniation and spilling of mtDNA and mtRNA into the cytoplasm. | Figure 4b and 4c |
| **Toxicity is from drug conjugate, not the peptide alone** | Cytotoxicity of (Fxr)<sub>3</sub> alone is > 100 uM in HeLa cells. | Horton *et al.* (2012)<sup>12</sup><br>• Figure 2b |
| **mtDox intercalates DNA** | mtDox fluorescent signal is quenched by the addition of DNA. | Chamberlain *et al.* (2013)<sup>15</sup><br>• Figure 2b |
| **mtDox poisons topoisomerase 2** | Inhibits TOP2 induced decatenation of kinetoplast DNA (kDNA). | Chamberlain *et al.* (2013)<sup>15</sup><br>• Figure 2b |
| **DNA damage is specific to mtDNA** | Comparative PCR inhibition in nDNA and mtDNA. | Chamberlain *et al.* (2013)<sup>15</sup><br>• Figure 2c |
| **mtDox avoids Dox-resistance** | Toxicity of mtDox unchanged in Dox-resistant A2780 cell line. | Chamberlain *et al.* (2013)<sup>15</sup><br>• Figure 3 |
| **mtDox causes an increase in mitobiogenesis markers** | Mitobiogenesis markers are increased following mtDox treatment including: SDHA, COX1, NRF1, and TFAM. | Jean, S. R. *et al.* (2015)<sup>16</sup><br>• Figure 5 |

## Methods

### Cell lines and culture methods

HCT116 *TP53* (-/-) cell line was acquired from Horizon Discovery (HD 104-001) and cultured in RPMI-1640 medium (Wisent) supplemented with 1 mM pyruvate (Gibco) and 5% or 10% FBS for drug treatment and cell passage respectively. Cells were also treated with Pen-Strep (100 U/mL) constitutively. Cells were incubated at 37 °C with 5% CO_2_. The doubling time of the cells was determined to be around 24 hours, with number of cell doublings represented by T*n*.

### Compounds

Doxorubicin hydrochloride salt (purity >99%) (LC Laboratories) was dissolved in DMSO (Sigma-Aldrich) and concentration was measured by absorbance at a wavelength of 488 nm with an extinction coefficient of 11,500 M^-1^ cm^-1^. mtDox was prepared as described previously^14^ and measured using the doxorubicin extinction coefficient. The LD_20_ for doxorubicin and mtDox were calculated at screen scale by plating 3 million cells per 150 mm plate, allowing cells to adhere overnight. Treatment was conducted by replacing media with a range of concentrations of drug-containing media. Drug effectiveness was calculated 72 hours later by washing cells with PBS, then lifting with trypsin and counting a representative population with a Countess II Automated Cell Counter counting only live cells as determined by trypan blue staining. The concentration of drug used was adjusted throughout the screen to maintain around 80% cell survival.

### Confocal microscopy localization

Microscopy was performed by treating HCT116 *TP53* (-/-) cells in an 8 Well Chamber µ-Slide (ibidi) for 2 hours with 8 μM Dox or mtDox, before addition of 150 nM MitoTracker Deep Red 30 mins before fixing cells with 4% formaldehyde then staining with NucBlue™ Fixed Cell ReadyProbes™ Reagent (DAPI) (Thermo Fisher). For WRNIP1-FLAG localization, the cells containing MitoTracker Deep Red were fixed then permeabilized with 0.2% Triton X-100 in PBS for 20 mins. The cells were blocked with 5% BSA in 0.1% Triton X-100 for 2 hours and the same solution was used for overnight primary antibody dilution at 4°C with anti-FLAG M2 (Sigma, F1804) 1:300. The cells were washed with PBS and stained with secondary goat anti-Mouse IgG (H+L) Highly Cross-Adsorbed Secondary Antibody, Alexa Fluor™ Plus 594 (Thermo Fisher, A-11005) 1:1000 for 1 hour at room temperature.

For WRNIP1-FLAG microscopy localization, HCT116 TP53 (-/-) cells were infected with lentivirus containing pGenLenti-CMV-WRNIP1-FLAG (ORF NM_020135.3). The construct was purchased from GenScript then packaged in-house using HEK293T cells with psPAX2, pMD2.G, and polyethylenimine (PEI) transfection reagent (M.W. ∼25000 linear, Alfa Aesar) in OptiMEM low serum media (Gibco) according to Addgene protocol for Lentivirus Production. Virus was collected after 72 hours. Selection of cells with successful incorporation of the construct was performed using 2 μg/mL puromycin 48 hours after transduction. Expression was verified by Western blotting using anti-FLAG M2 (Sigma, F1804) 1:1000 and anti-WRNIP1 (Santa Cruz, sc-377402). The cells were stained with MitoTracker Deep Red (200 nM) for 30 mins then fixed with 4% paraformaldehyde in 8 Well Chamber µ-Slides (ibidi). Cells were permeabilized with 0.2% Triton X-100 in PBS for 20 mins. The cells were blocked with 5% BSA in 0.1% Triton X-100 for 2 hours and the same solution was used for overnight primary antibody dilution at 4°C with anti-FLAG M2 (Sigma, F1804) 1:300. The cells were washed with PBS and stained with secondary goat anti-Mouse IgG (H+L) Highly Cross-Adsorbed Secondary Antibody, Alexa Fluor™ Plus 594 (1:1000) for 1 hour at room temperature.

### Lentiviral library MOI determination

The TKOv3 library was a gift from the Anger’s Lab (Addgene 90294) prepared in the all-in-one lentiCRISPRv2 backbone. Performed viral titration by transducing HCT116 *TP53* (-/-) cells (50,000 cells per 6-well plate) with increasing volume of virus in duplicate using 5 μg/mL polybrene. One of each duplicate was treated with 2 μg/mL puromycin (Sigma) after 24 hours while the other received fresh media. Cells we incubated for another 48 hours then fixed with MeOH and stained with crystal violet. The stain was released with 10% acetic acid in water and measured at 590 nm absorbance. MOI was determined by dividing puromycin-treated by puromycin-untreated for each virus volume and generating a curve to calculate which volume would give an MOI of 0.3 for screen infection. All volumes and cell counts for screen infection were scaled up by dish surface area.

### CRISPR/Cas9 screen

HCT116 *TP53* (-/-) were infected in a pool such that accounting for an estimated MOI of 0.3, there would be enough surviving cells to represent at least 200-fold library coverage which translates to a minimum of around 50 million cells plated (T0). Cells were plated in 150 mm dishes at 4 million cells per plate and transduced with a pooled medium containing library virus, 1 mM pyruvate, 5 μg/mL polybrene and 100 U/mL Pen-Strep. One plate was left without virus to calculate the true screen MOI. Media was replaced 24 hours later with media containing 2 μg/mL puromycin and selection took place over an additional 48 hours (T1). Cells were then split into new 150 mm dishes based on their intended treatment condition, maintaining 200-fold library coverage for each condition (T4). For this screen, 15 million cells were plated per condition in triplicate, at 3 million cells per 150 mm dish. On T5, the cells were treated with their respective condition: DMSO, Dox (LD_20_), or mtDox (LD_20_) and left to incubate for 72 hours before a new passage. With each passage 200-fold coverage was maintained, while the remaining cells were frozen in pellets or used for flow cytometry experiments. A cycle of passage and treatment was conducted until T20 when the final pellets were collected. The remaining cells were plated again in order to grow enough cells for mtDNA sequencing.

### Genomic screen DNA extraction and sequencing

Genomic DNA was extracted from screen pellets using QIAamp DNA Blood Maxi Kit (QIAGEN) and gRNAs were isolated using PCR, followed by a second PCR reaction with Illumina TruSeq i5 and i7 indices. The protocol was adapted from MacLeod *et al.*^52^.

### Screen analysis

Sequencing data was mapped to library guides and converted to read count using the MAGeCK count algorithm^24^. Read count was analyzed using the DrugZ algorithm comparing T20 drug-treated conditions to T20 DMSO control^25^. Genes are ranked based on a normalized Z-score (normZ) producing a list of susceptibility and resistance genes in each drug context. Full data for drugZ analysis provided in Source Data. Includes source data for Figures 1 c & d, and Figure 2a.

### CRISPR knockout generation

Guides were designed separate from the library using the BROAD Institute sgRNA designer tool. For validation experiments, guides were cloned into a lentiCRISPRv2 vector for lentiviral knockout. Some sgRNA plasmids were ordered through GenScript custom cloning or through their catalog. Virus was generated in HEK293T cells using psPAX2, pMD2.G, and polyethylenimine (PEI) transfection reagent in OptiMEM low serum media (Gibco) according to addgene protocol. Virus was collected after 24 and 48 hours. Viruses were used to transduce HCT116 *TP53* (-/-) cells which were selected for lentiCRISPRv2 incorporation using 2 μg/mL puromycin after 48 hours. Knockouts were verified by Western blotting or TIDE sequencing (Table S1).

For the stable scarless knockout of WRNIP1, the px459 plasmid cloned with sgWRNIP1 was transfected directly to HCT116 *TP53* (-/-) cells using PEI as described above. Cells were treated with puromycin for 48 hours and surviving clones were picked and grown in monoclonal populations until knockout could be verified using TIDE sequencing.

### Clonogenic validation experiments

Cell lines were collected, resuspended thoroughly, and plated at low density (250 cells per well of a 6-well plate). Cells were left to adhere over 48 hours before treatment with Dox or mtDox. Cells were grown for an additional 12 days without removing drug until visible colonies could be counted. Cells were immediately fixed with ice-cold MeOH and left in the freezer for at least 24 hours before staining with crystal violet and washing away excess stain. Images of the plates were taken with BioRad ChemiDoc Imaging System then processed with ImageJ to count total pixel area coverage. Area counts were normalized to vehicle control to correct for differences in cell seeding and growth rate between knockouts. Relative survival of each target of interest was compared to survival in a sgLacZ negative control experiment plated and treated on the same days using same stock of compound.

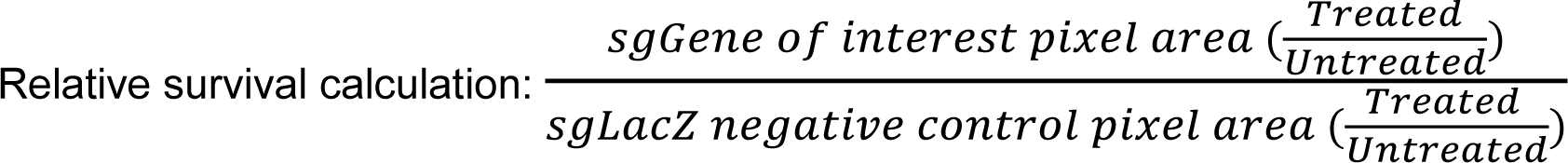

For the short-term 72-hour survivability treatment as measured by PrestoBlue (Thermo Fisher), the cells were plated in 12-well plates at 7500 cells per well. The media was replaced with mtDox treatment media 24 hours later and the cells were left to grow for an additional 72 hours. Media was replaced with 300 μL fresh media containing 10x diluted PrestoBlue and incubated for 90 mins before 200 μL from each well was transferred to a Corning Costar® 96-Well Black Polystyrene Plate. The fluorescence was read from the 96-well plate using a plate reader set for Ex/Em of 560/590 nm.

### Mitochondrial DNA isolation, deep sequencing, and analysis

To isolate mtDNA for sequencing, 150-200 million cells were collected and washed with PBS before resuspension in Buffer A with protease inhibitors. Cells were left to swell for 5 minutes before adjusting sucrose concentration to 250 mM and using a dounce homogenizer to break cell membrane. Lysed cells were spun for 3 mins at 1300 g then the supernatant was collected and spun at 15,000g for 15 mins and washed 3 times with 250 mM sucrose in T_10_E_20_ buffer. A sucrose gradient was set up with 1.7 M sucrose on the bottom layer and 1.0 M sucrose on the top layer. The mitochondrial pellet was set on top of the gradient and spun in a Beckman Coulter Optima MAX-XP ultracentrifuge for 40 mins at 75,000g using a swinging bucket rotor. Mitochondria were collected from the interface between 1.7 and 1.0 M sucrose and washed 3 times with 250 mM sucrose in T_10_E_20_ at 15,000g for 15 mins. mtDNA was extracted from the resultant pellet using GenElute Mammalian Genomic DNA Miniprep Kit (Sigma). The remaining nDNA contamination was removed by treatment with Plasmid-Safe ATP-Dependent DNase (Lucigen) to digest linear DNA^53^. Enrichment of mtDNA was verified using qPCR and primers targeting MT-ND2 and nuclear PCNA. Library prep and sequencing was performed by the Donnelly Sequencing Centre using the Nextera XT library prep then sequenced using single-end Illumina MiSeq v2 Micro flow cell. The analysis was prepared using bowtie2-build to generate a consensus mtDNA sequence from NC_012920 as a reference for bowtie2 to assign sequencing reads as belonging to mitochondria^50^. The alignments were processed through samtools, followed by variant calling using freebayes and bcftools^51^. The number of mutations was normalized to the number of successful read alignments to account for differing efficiencies in mtDNA enrichment and sequencing.

### mtDNA copy number analysis

HCT116 *TP53* (-/-) cells were treated with mtDox and collected at various time points post-treatment using trypsin. Whole cell DNA was extracted from pellets using QIAGEN DNeasy Blood & Tissue Kit. Relative quantification of the mtDNA copy number was performed using SYBR-green based qPCR for mtDNA specific primers with an 18S endogenous nuclear control (Supplementary Table 2). A standard protocol was used for Applied Biosystems™ Power SYBR™ Green PCR Master Mix on a BioRad CFX384 Real-Time PCR System.

### Flow cytometry for JC-1, MitoTracker Deep Red, and H2DCFDA

HCT116 *TP53*(-/-) cells were plated at 25,000 cells per well of a 12-well plate and allowed to adhere for 24 hours before treatments began. MitoTracker or JC-1 stain was added 30 minutes before collection of cells using trypsin. Cells were washed with PBS before running through a Beckman CytoFLEX S flow cytometer using the APC channel for MitoTracker Deep Red, and FITC (monomers) and PE (aggregates) for JC-1 dye. Cells were gated for forward and side scattering. JC-1 summary values were calculated by dividing PE by FITC mean fluorescence. ROS probe H2DCFDA was prepared fresh each time and 10 μM was added to cells post-trypsin collection in PBS for 20 mins before washing with PBS and running through a Beckman CytoFLEX S flow cytometer using the FITC channel and gating for forward and side scattering (Provided in Source Data).

### Proteinase K

The isolation and digestion of mitochondria was performed as described previously^17^ using HCT116 *TP53*(-/-) cells with the following alterations: only 50 μg of mitochondrial isolate was used for digestion, and upon the addition of 2 volumes of ice-cold 20% TCA the protein was incubated on ice for 30 mins, the precipitate was spun for 5 mins at 12000 g before washing with 200 μL ice cold acetone then spun down again before removal of supernatant and drying with cap off at 95 °C for 1 min. Pellets were stored at -20 °C. For use in a Western blot, the pellet was resuspended in 2x Laemmli buffer with β-mercapto ethanol (Bio-Rad), sonicated to dissolve pellet, and split over 2 wells.

### Seahorse Mito Stress Test

Followed suggested protocol from Agilent with the following modifications. HCT116 *TP53*(-/-) WT and WRNIP1 monoclonal knockout (KO) cells were plated in a Seahorse cell plate 24 hours prior to experiment. After the cells adhered to the plate, the media was switched with media containing 8 μM mtDox or DMSO. After 6 hours treatment, the protocol proceeded exactly as suggested by Agilent. Cells were normalized to cell count using a Hoechst stain and a Cytation 5 Cell Imaging Multimode Reader.

### Co-immunoprecipitation

HCT116 *TP53*(-/-) cells were transfected with a FLAG-tagged RIG-I expression vector purchased from Origene (RC217615) using Lipofectamine 3000 in six 150 mm dishes plated to 90% confluency the day before. Cells were grown for an additional 72 hours before collection by trypsin, amalgamation of all plates, then equal redistribution of a homogenous transfected cell population to new 150 mm plates. The co-immunoprecipitation was conducted the following day using 8 μM mtDox or vehicle control. Each time point was collected via cell scraping from an individual 150 mm dish according to the protocol described by Tan *et al.* using RIPA lysis buffer (CST, 9806S) supplemented with protease inhibitor cocktail (CST, 5872S) and 1 mM PMSF^35^. The FLAG-tagged RIG-I was pulled down using anti-FLAG M2 magnetic beads purchased from Sigma-Aldrich (M8823). The proteins were eluted from beads in 30 μL of 4x Laemmli buffer heated to 95 °C for 5 mins and the entire elution was run on a Western blot using anti-FLAG (Sigma-Aldrich F3165) and anti-WRNIP1 (Abcam, ab4731 or Santa Cruz, sc-377402).

### Western blotting

In general, HCT116 *TP53*(-/-) cells were collected through trypsinization followed by a PBS wash, then RIPA lysis of the pellets for 30 minutes at 4°C with agitation. The lysis buffer was freshly supplemented with protease inhibitor cocktail (CST, 5872S) and 1 mM PMSF. The lysate was then spun at 15,000g for 15 mins at 4°C to eliminate the insoluble fraction. In the case of the p-IRF3 Western blot, cells were collected directly from 6-well plates via scraping using cold RIPA buffer supplemented with protease and phosphatase inhibitors (CST, 5872S) and 1 mM PMSF. The spun down lysates were measured for total protein using a BCA assay and 15 μg of protein was diluted in 2x Laemmli buffer containing β-mercapto ethanol (Bio-Rad) and heated to 95 °C for 5 mins. Boiled lysates were loaded on a 4-15% Tris-glycine gradient gel and run at 150 V using Tris/Glycine/SDS buffer (Bio-Rad, 1610732) until loading buffer reached the bottom of the gel. The gel was transferred to a PVDF membrane using Tris/Glycine buffer (Bio-Rad, 1610734) at 80 V for 75 mins. Membranes were blocked with 5% skim milk in TBST for an hour at room temperature. The p-IRF3 Western blots were blocked with 5% BSA in TBST. Primary antibodies were incubated on membrane in blocking buffer overnight at 4°C with agitation. Secondary antibodies were incubated for 1 hour in TBST at room temperature with agitation. TBST was used to wash membrane 4 times for 5 mins each between every step. Primary antibodies and dilutions used were: anti-WRNIP1 (Abcam, ab4731 or Santa Cruz, sc-377402) 1:500; anti-FLAG M2 (Sigma-Aldrich F3165) 1:1000; anti-BCL-XL (Abcam, ab32370) 1:500; anti-TFAM (Abcam, ab131607) 1:500; anti-β-tubulin (CST, 2128S) 1:1000; anti-GAPDH (CST, 2118S) 1:2000; anti-p-IRF3 (CST, 4947S) 1:1000; anti-MTCO2 (Abcam, ab91317) 1:500; anti-Lamin B (Abcam, ab16048) 1:1000. Secondary antibodies used were anti-rabbit or anti-mouse IgG, HRP-linked antibody (CST, 7074S, 7076S). Blots were developed using SuperSignal West Pico PLUS Chemiluminescent Substrate and imaged using BioRad ChemiDoc Imaging System.

### Cytoplasmic mtDNA/mtRNA release

Digitonin-based protocol adapted from References^43,44,54^. HCT116 *TP53*(-/-) cells were treated with 8 μM mtDox for up to 6 hours. For both mtDNA and mtRNA release, the cell pellet was resuspended in 40 uL PBS, then 20 uL each was used to represent the cytoplasmic fraction which underwent digitonin isolation, while the other 20 uL was used to normalize cell number (WC). For mtDNA measurements, an mtDNA-specific primer was used with the cytoplasmic fractions and an 18S rDNA primer was used on WC. Undiluted cytoplasmic fraction was measured while the WC fractions were diluted 5x. For the mtRNA measurements, both the cytoplasmic and WC fractions were reverse transcribed using iScript cDNA Synthesis Kit (Bio-Rad) and 12S mitochondrial rRNA primer was used for both the cytoplasmic and WC fractions (Supplementary Table 2). qPCR was performed using Applied Biosystems™ Power SYBR™ Green PCR Master Mix on a BioRad CFX384 Real-Time PCR System.

### Comet assay

This assay was performed according to the specifications of Abcam’s comet assay kit (ab238544) using HCT116 *TP53*(-/-) with and without WRNIP1 monoclonal KO . The gels were run at 100 mV for 30 mins and the agarose-embedded cells were visualized using a confocal fluorescent microscope collecting up to 10 representative images per sample. The comet tails in each image were analyzed using the automated OpenComet plugin for ImageJ.

### yH2AX Microscopy

HCT116 *TP53*(-/-) cells were seeded at 10,000 cells per well in an Ibidi microslide and left to adhere for 48 hours. Cells were treated with either 8 uM mtDox or 25 nM Dox, with and without 10 mM *N*-acetyl cysteine (NAC). After 24 hours incubation, cells were fixed with 4% formaldehyde, permeabilized with 0.4% Triton X-100, and blocked with 5% FBS. Primary incubation was set up overnight at 4 °C with anti-γH2AX antibody (CST, 9718S) 1:400. Secondary anti-rabbit Alexa Fluor 488 (Thermo Fisher Scientific, A11008) 1:1000 was added for 1 hour at room temperature. Washed cells were mounted in 0.1 ug/mL DAPI. Imaged using Zeiss inverted confocal with 63x objective.

### Quantification and Statistical Analysis

Statistical analysis was performed using the GraphPad Prism 8 software using an unpaired Student’s *t*-test to calculate significance. A result was considered significant with a *p*-value < 0.05. Data is represented as mean ± s.d. The number of replicates for each experiment is defined in the figure captions, with n representing discrete biological replicates.

### Chemical Compounds

No new chemical compound was reported in this study, however the (Fxr)_3_ peptide and mtDox are synthesized in-house and are available upon request to the lead contact.

### Data Availability

The Excel file containing the results of the drugZ analysis for doxorubicin and mtDox is available upon request to the lead contact.

## Notes

### Competing Interest Statement

The authors have declared no competing interest.

## References

1. Friedman, J. R. & Nunnari, J. (2014) Mitochondrial form and function. Nature 505, 335–343.

2. Johnston, I. G. & Williams, B. P. (2016) Evolutionary inference across eukaryotes identifies specific pressures favoring mitochondrial gene retention. Cell Syst. 2, 101–111.

3. Rath, S., et al. (2021) MitoCarta3. 0: an updated mitochondrial proteome now with sub-organelle localization and pathway annotations. Nucleic Acids Res. 49, D1541–D1547.

4. Herzig, S. & Shaw, R. J. (2018) AMPK: guardian of metabolism and mitochondrial homeostasis. Nat. Rev. Mol. Cell Biol. 19, 121–135.

5. Kazak, L., Reyes, A. & Holt, I. J. (2012) Minimizing the damage: repair pathways keep mitochondrial DNA intact. Nat. Rev. Mol. Cell Biol. 13, 659–671.

6. Prakash, A. & Doublié, S. (2015) Base excision repair in the mitochondria. J. Cell. Biochem. 116, 1490–1499.

7. Copeland, W. C. & Longley, M. J. (2014) Mitochondrial genome maintenance in health and disease. DNA Repair 19, 190–198.

8. Mason, P. A., Matheson, E. C., Hall, A. G. & Lightowlers, R. N. (2003) Mismatch repair activity in mammalian mitochondria. Nucleic Acids Res. 31, 1052–1058.

9. Tadi, S. K., et al. (2016) Microhomology-mediated end joining is the principal mediator of double-strand break repair during mitochondrial DNA lesions. Mol. Biol. Cell 27, 223–235.

10. Moretton, A., et al. (2017) Selective mitochondrial DNA degradation following double-strand breaks. PLoS One 12, e0176795.

11. Wu, Z., Sainz, A. G. & Shadel, G. S. (2021) Mitochondrial DNA: cellular genotoxic stress sentinel. Trends Biochem. Sci. 46, 812–821.

12. Horton, K. L., Pereira, M. P., Stewart, K. M., Fonseca, S. B. & Kelley, S. O. (2012) Tuning the Activity of Mitochondria-Penetrating Peptides for Delivery or Disruption. ChemBioChem 13, 476–485.

13. Stewart, K. M., Horton, K. L. & Kelley, S. O. (2008) Cell-penetrating peptides as delivery vehicles for biology and medicine. Org. Biomol. Chem. 6, 2242–2255.

14. Capranico, G., Kohn, K. W. & Pommier, Y. (1990) Local sequence requirements for DNA cleavage by mammalian topoisomerase II in the presence of doxorubicin. Nucleic Acids Res. 18, 6611–6619.

15. Chamberlain, G. R., Tulumello, D. V. & Kelley, S. O. (2013) Targeted delivery of doxorubicin to mitochondria. ACS Chem. Biol. 8, 1389–1395.

16. Jean, S. R., et al. (2015) Mitochondrial targeting of doxorubicin eliminates nuclear effects associated with cardiotoxicity. ACS Chem. Biol. 10, 2007–2015.

17. Buondonno, I., et al. (2016) Mitochondria-targeted doxorubicin: a new therapeutic strategy against doxorubicin-resistant osteosarcoma. Mol. Cancer Ther. 15, 2640–2652.

18. Wisnovsky, S., Jean, S. R., Liyanage, S., Schimmer, A. & Kelley, S. O. (2016) Mitochondrial DNA repair and replication proteins revealed by targeted chemical probes. Nat. Chem. Biol. 12, 567.

19. Wisnovsky, S., Sack, T., Pagliarini, D. J., Laposa, R. R. & Kelley, S. O. (2018) DNA polymerase θ increases mutational rates in mitochondrial DNA. ACS Chem. Biol. 13, 900–908.

20. Hart, T., et al. (2017) Evaluation and design of genome-wide CRISPR/SpCas9 knockout screens. G3: Genes Genomes Genet. 7, 2719–2727.

21. Hart, T., et al. (2015) High-resolution CRISPR screens reveal fitness genes and genotype-specific cancer liabilities. Cell 163, 1515–1526.

22. Geisinger, J. M. & Stearns, T. (2020) CRISPR/Cas9 treatment causes extended TP53-dependent cell cycle arrest in human cells. Nucleic Acids Res. 48, 9067–9081.

23. Olivieri, M., et al. (2020) A genetic map of the response to DNA damage in human cells. Cell 182, 481–496. e21.

24. Li, W., et al. (2014) MAGeCK enables robust identification of essential genes from genome-scale CRISPR/Cas9 knockout screens. Genome Biol. 15, 554.

25. Colic, M., et al. (2019) Identifying chemogenetic interactions from CRISPR screens with drugZ. Genome Med. 11, 1–12.

26. Sampson, A., Peterson, B. G., Tan, K. W. & Iram, S. H. (2019) Doxorubicin as a fluorescent reporter identifies novel MRP1 (ABCC1) inhibitors missed by calcein-based high content screening of anticancer agents. Biomed. Pharmacother. 118, 109289.

27. Schellenberg, M. J., et al. (2017) ZATT (ZNF451)–mediated resolution of topoisomerase 2 DNA-protein cross-links. Science 357, 1412–1416.

28. Schaupp, C. M., White, C. C., Merrill, G. F. & Kavanagh, T. J. (2015) Metabolism of doxorubicin to the cardiotoxic metabolite doxorubicinol is increased in a mouse model of chronic glutathione deficiency: A potential role for carbonyl reductase 3. Chem. Biol. Interact. 234, 154–161.

29. Lyu, Y. L., et al. (2007) Topoisomerase IIβ–mediated DNA double-strand breaks: implications in doxorubicin cardiotoxicity and prevention by dexrazoxane. Cancer Res. 67, 8839–8846.

30. Nordgren, K. K. & Wallace, K. B. (2014) Keap1 redox-dependent regulation of doxorubicin-induced oxidative stress response in cardiac myoblasts. Toxicol. Appl. Pharmacol. 274, 107–116.

31. Matsumaru, D. & Motohashi, H. (2021) The KEAP1-NRF2 system in healthy aging and longevity. Antioxidants 10, 1929.

32. Wang, X., et al. (2008) Nrf2 enhances resistance of cancer cells to chemotherapeutic drugs, the dark side of Nrf2. Carcinogenesis 29, 1235–1243.

33. Mi, H., et al. (2021) PANTHER version 16: a revised family classification, tree-based classification tool, enhancer regions and extensive API. Nucleic Acids Res. 49, D394–D403.

34. Huang, S., et al. (2018) Mitochondrial tyrosyl-DNA phosphodiesterase 2 and its TDP2 short isoform. EMBO Rep. 19, e42139.

35. Yoshimura, A., Seki, M. & Enomoto, T. (2017) The role of WRNIP1 in genome maintenance. Cell Cycle 16, 515–521.

36. Socha, A., et al. (2020) WRNIP1 Is Recruited to DNA Interstrand Crosslinks and Promotes Repair. Cell Rep. 32, 107850.

37. Tirman, S., Cybulla, E., Quinet, A., Meroni, A. & Vindigni, A. (2021) PRIMPOL ready, set, reprime! Crit. Rev. Biochem. Mol. Biol. 56, 17–30.

38. Tan, P., et al. (2017) Assembly of the WHIP-TRIM14-PPP6C mitochondrial complex promotes RIG-I-mediated antiviral signaling. Mol. Cell 68, 293–307. e5.

39. Li, M., Wang, D., He, J., Chen, L. & Li, H. (2020) Bcl-xl: A multifunctional anti-apoptotic protein. Pharmacol. Res. 151, 104547.

40. Murphy, M. P. & Hartley, R. C. (2018) Mitochondria as a therapeutic target for common pathologies. Nat. Rev. Drug Discov. 17, 865–886.

41. Grazioli, S. & Pugin, J. (2018) Mitochondrial damage-associated molecular patterns: from inflammatory signaling to human diseases. Front. immunol. 9, 832.

42. Pérez-Treviño, P., Velásquez, M. & García, N. (2020) Mechanisms of mitochondrial DNA escape and its relationship with different metabolic diseases. Biochim. Biophys. Acta - Mol. Basis Dis. 1866, 165761.

43. Tigano, M., Vargas, D. C., Tremblay-Belzile, S., Fu, Y. & Sfeir, A. (2021) Nuclear sensing of breaks in mitochondrial DNA enhances immune surveillance. Nature 591, 477–481.

44. Wu, Z., et al. (2019) Mitochondrial DNA stress signalling protects the nuclear genome. Nat. Metab. 1, 1209–1218.

45. Strano, S. et al. (2007) Mutant p53: an oncogenic transcription factor. Oncogene 26, 2212–2219.

46. Marei, H. E. et al. (2021) p53 signaling in cancer progression and therapy. Cancer Cell Int. 21, 1–15.

47. Park, J., Zhuang, J., Li, J. & Hwang, P. M. (2016) p53 as guardian of the mitochondrial genome. FEBS Lett. 590, 924–934.

48. Follis, A. V. et al. (2013) PUMA binding induces partial unfolding within BCL-xL to disrupt p53 binding and promote apoptosis. Nat. Chem. Biol. 9, 163–168.

49. Zhang, W. et al. (2020) The mitochondrial protein MAVS stabilizes p53 to suppress tumorigenesis. Cell Rep. 30, 725–738. e4.

50. Langmead, B. & Salzberg, S. L. (2012) Fast gapped-read alignment with Bowtie 2. Nature methods 9, 357.

51. Li, H. (2011) A statistical framework for SNP calling, mutation discovery, association mapping and population genetical parameter estimation from sequencing data. Bioinformatics 27, 2987–2993.

52. MacLeod, G., et al. (2019). Genome-Wide CRISPR-Cas9 Screens Expose Genetic Vulnerabilities and Mechanisms of Temozolomide Sensitivity in Glioblastoma Stem Cells. Cell reports 27, 971–986.

53. Gould, M. P., et al. (2015) PCR-free enrichment of mitochondrial DNA from human blood and cell lines for high quality next-generation DNA sequencing. PLoS One 10, e0139253.

54. West, A. P., Shadel, G. S. & Ghosh, S. (2011) Mitochondria in innate immune responses. Nat. Rev. Immunol. 11, 389–402.

