## Supplementary Figures and Tables for "CRISPR Screening in Tandem with Targeted mtDNA Damage Reveals WRNIP1 Essentiality"

[1]Department of Pharmaceutical Sciences, Leslie Dan Faculty of Pharmacy, University of Toronto, Toronto, Ontario M5S 3M2, Canada. [2]Department of Biochemistry, Faculty of Medicine, University of Toronto, Toronto, Ontario M5S 1A8, Canada. [3]Terrence Donnelly Centre for Cellular & Biomolecular Research, University of Toronto, Toronto, Ontario M5S 3E1, Canada. [4]Department of Chemistry, Faculty of Arts and Science, University of Toronto, Toronto, Ontario M5S 3H6, Canada. [5]Department of Chemistry, Weinberg College of Arts & Sciences, Northwestern, Illinois 60208, United States.

#### Table S1: sgRNA Sequences and TIDE Primers

| Gene Target | sgRNA | TIDE_Fwd | TIDE_Rev |
|---|---|---|---|
| WRNIP1 – T1 | GCAGTCCGTTCAACCCAGCT | GGTGAGAATAAGAGGCAAAGGC | TGAGGCACGCTAAAGCCAAA |
| WRNIP1 – T2 | GGCATCATCTGCTGGCACAC | AACGGCCACGAACTACACTT | GCATCTGTCGGATCTCCTCG |
| MCM9 | CCAAGATCTGTGCTGACCAC | ACATAGGCCTGAACATCTGGA | GCAATTCACTTTCAAACTGCCC |
| RECQL5 | GGATGAAGCTGCCATCTCTG | CTGGGAAATCCTCCAGGGAC | GCCTGGCTGAGAAAGCTAGAA |
| TDP2 | GAACTCCTACTTCGAGCCTC | AAGATGGAGTTGGGGAGTTGC | CAGGAAGCAAGGAGACTTCCA |
| XRCC4 | TTTGTTATTACACTTACTGA | TGTGTAGCTGAGAGGCCAGT | AGCCAAGATCCTAAATGTGGGT |
| RMI2 | AGCGGGGACTTCTCGGTCCG | AGCCAAGATCCTAAATGTGGGT | AGGAGAAGTACAGGAGGGTCTC |
| PSMC3IP | AATCGTGGCCCTCACTGCTA | GCCGCTCCTTAACTGAAAACAA | AGCTCTGAAACCACATCCGC |
| MND1 | GTTGGAGGTTCTGGAATCTC | Appropriate primer pair not found | |
| C17orf53 | GCCCACTCACCAGACAACCT | ACTTGGAGACCTGGGAAGTCA | AGGGAAGGTCAGTGTCCTGAT |

#### Table S2: Sequences Used for qPCR

| Target | Fwd | Rev |
|---|---|---|
| mtDNA Purification Test | | |
| mtDNA marker (MT-ND2) | CCCCACAAACCCCATTACTAAACCCA | TTTCATCATGCGGAGATGTTGGATGG |
| nDNA marker (PCNA) | GCCACTCCACTCTCTTCAACG | CAACTGAAAGACAGGAAGATGGTT |
| mtDNA Copy Number and Cytoplasmic Release | | |
| mtDNA | GCCCCCGATATGGCGTTTCC | GTTCAACCTGTTCCTGCTCC |
| 18S | TAGAGGGACAAGTGGCGTTC | CGCTGAGCCAGTCAGTGT |
| mtRNA Cytoplasmic Release | | |
| mtRNA (12S) | TAGCCCTAAACCTCAACAGT | TGCGCTTACTTTGTAGCCTTCAT |

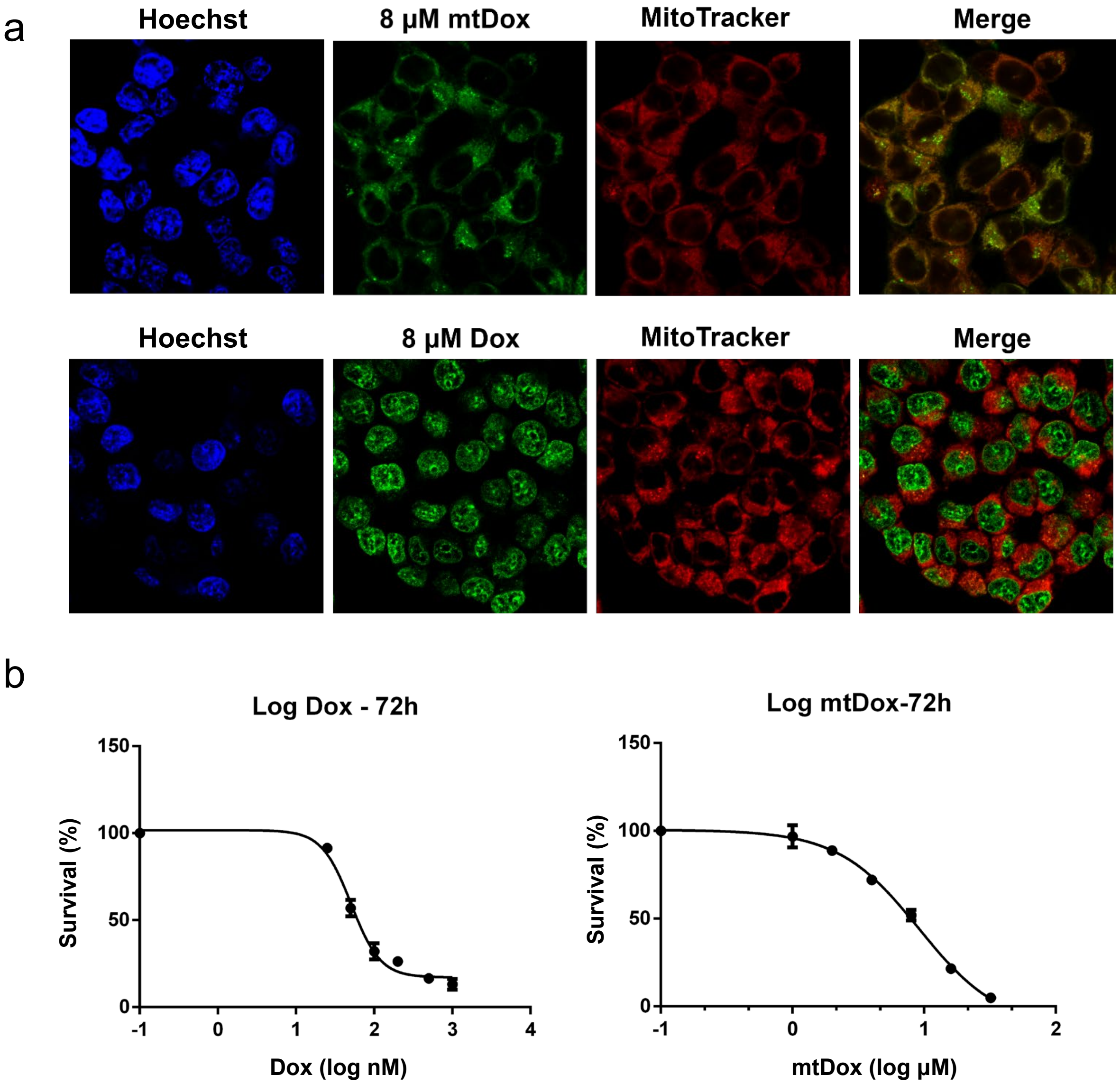


**Figure S1: Differential localization and potency of mtDox versus parent compound Dox. (a)** All probes localized within an hour when cells were fixed using 4% paraformaldehyde and visualized using a confocal microscope. mtDox and Dox are visible using the 488 channel. MitoTracker Deep Red was visualized using 633 channel. **(b)** Dose-response curves for Dox and mtDox after 72 hours of treatment. Concentrations tested for Dox were: 0, 25, 50, 100, 200, 500, and 1000 nM. Concentrations for mtDox tested were: 0, 1, 2, 4, 8, 16, and 32 μM. $LD_{20}$ values for Dox and mtDox were calculated as 30 nM, and 3.5 μM, respectively. $LD_{80}$ values for Dox and mtDox were calculated as 100 nM and 20 μM, respectively. Survival measured as cell count lifted after 72 hours treatment with indicated compound in 150 mm dishes.

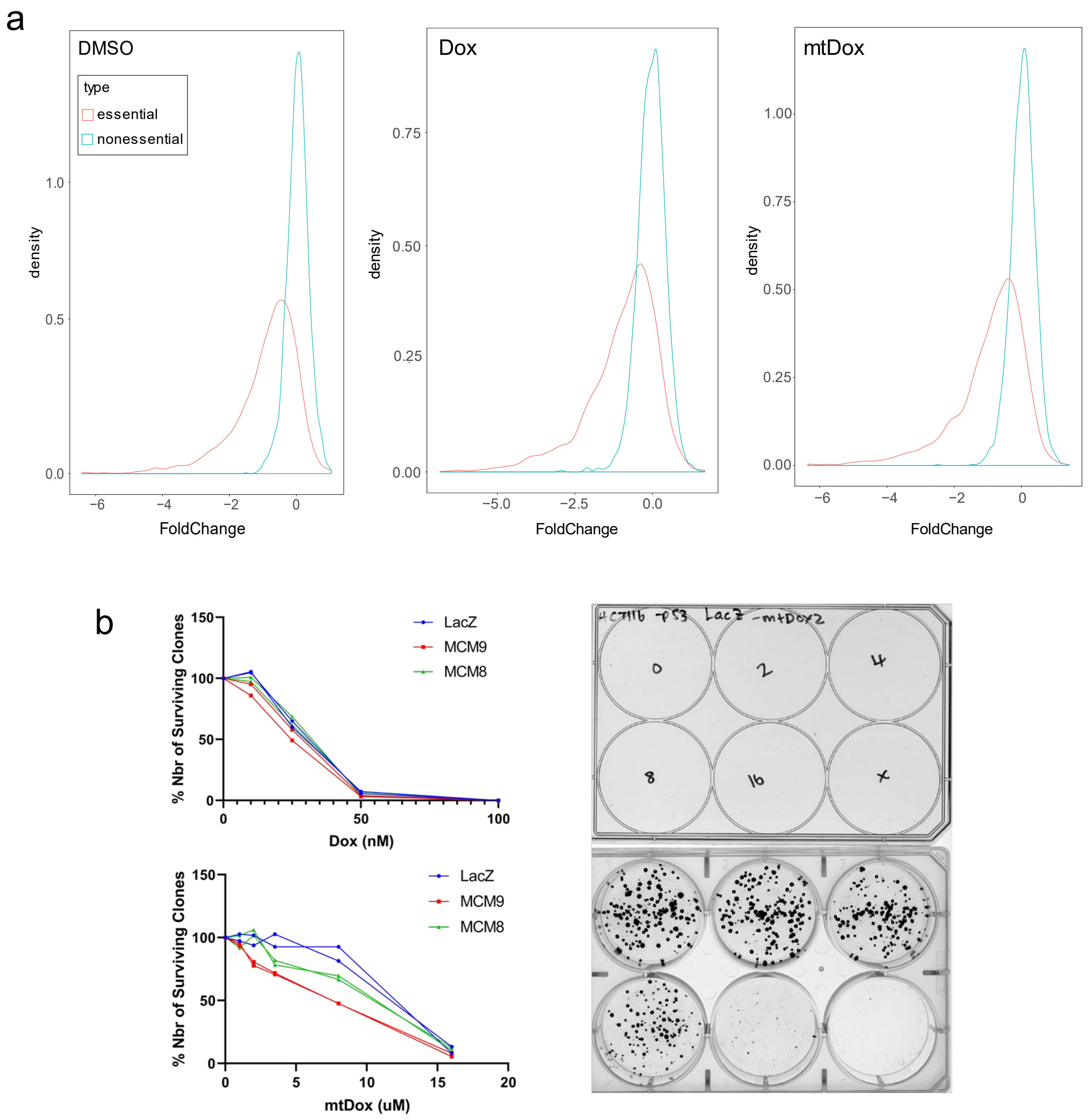


**Figure S2: Quality control analysis and validation of screen results. (a)** If screens were conducted according to expected experimental outcome, should see increased dropout of essential genes and little dropout of non-essential genes. All conditions screened displayed this phenotype. **(b)** Representative analysis of clonogenic survival assays used for screen validation.

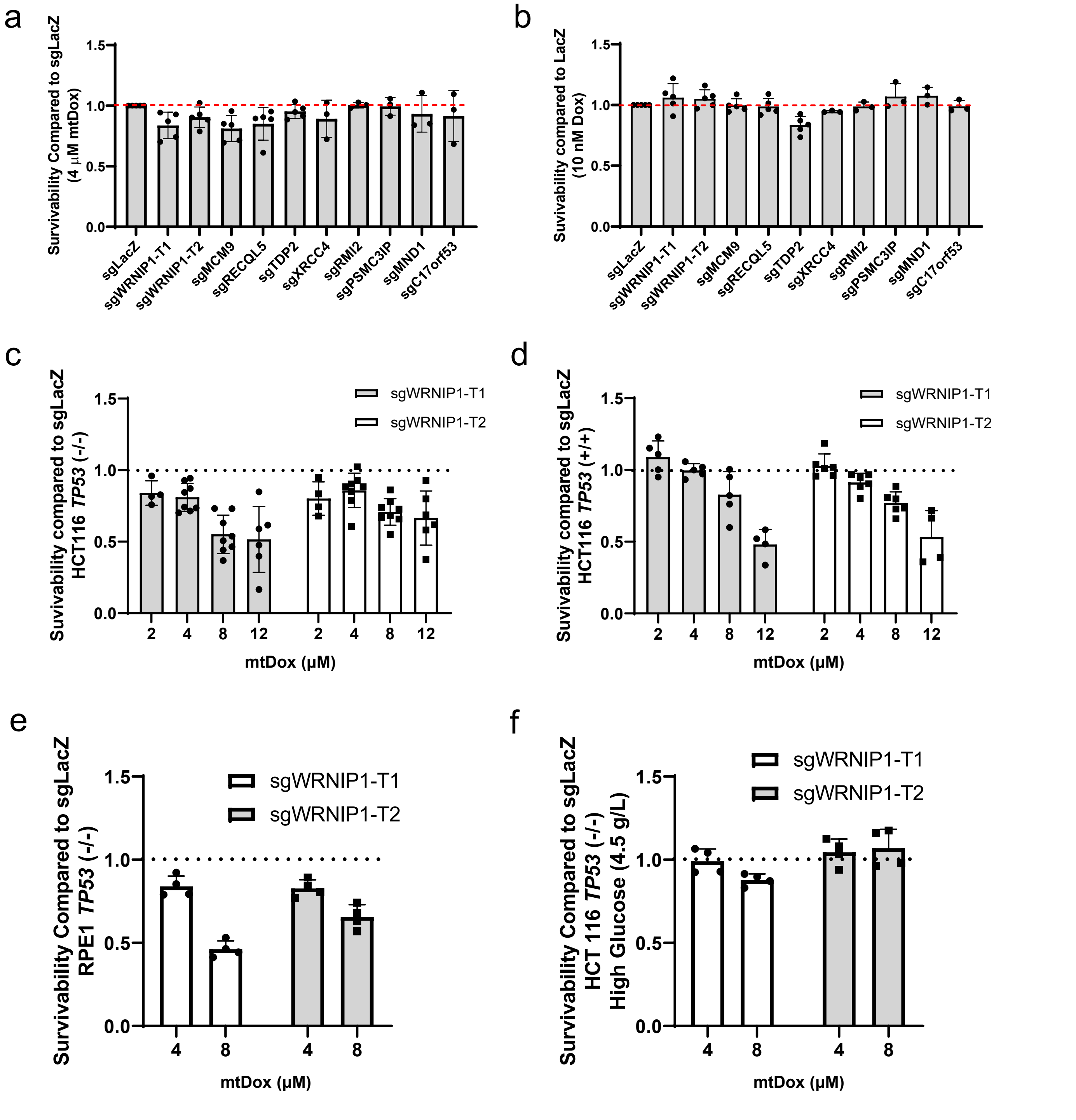


**Figure S3: Clonogenic analysis of differential cell line and growth conditions.**
**(a)** Survivability was also tested at 4 μM mtDox but displays less differentiation between KOs compared to 8 μM mtDox. **(b)** Survivability was also tested at 10 nM Dox, however differential survivability of positive control sgTDP2 is less significant than at 25 nM. **(c)** Alternative display of all clonogenic concentrations tested for HCT116 *TP53* (-/-). **(d)** Clonogenic survivability in HCT116 *TP53*(+/+) in a range of concentrations. **(e)** Clonogenic survivability in alternate cell line RPE1 *TP53* (-/-) representing a cell line lacking p53 but with non-truncated TFAM. **(f)** Clonogenic survivability of CRISPR/Cas9 cell line HCT116 *TP53* (-/-) with high D-glucose concentrations (4.5 g/L).

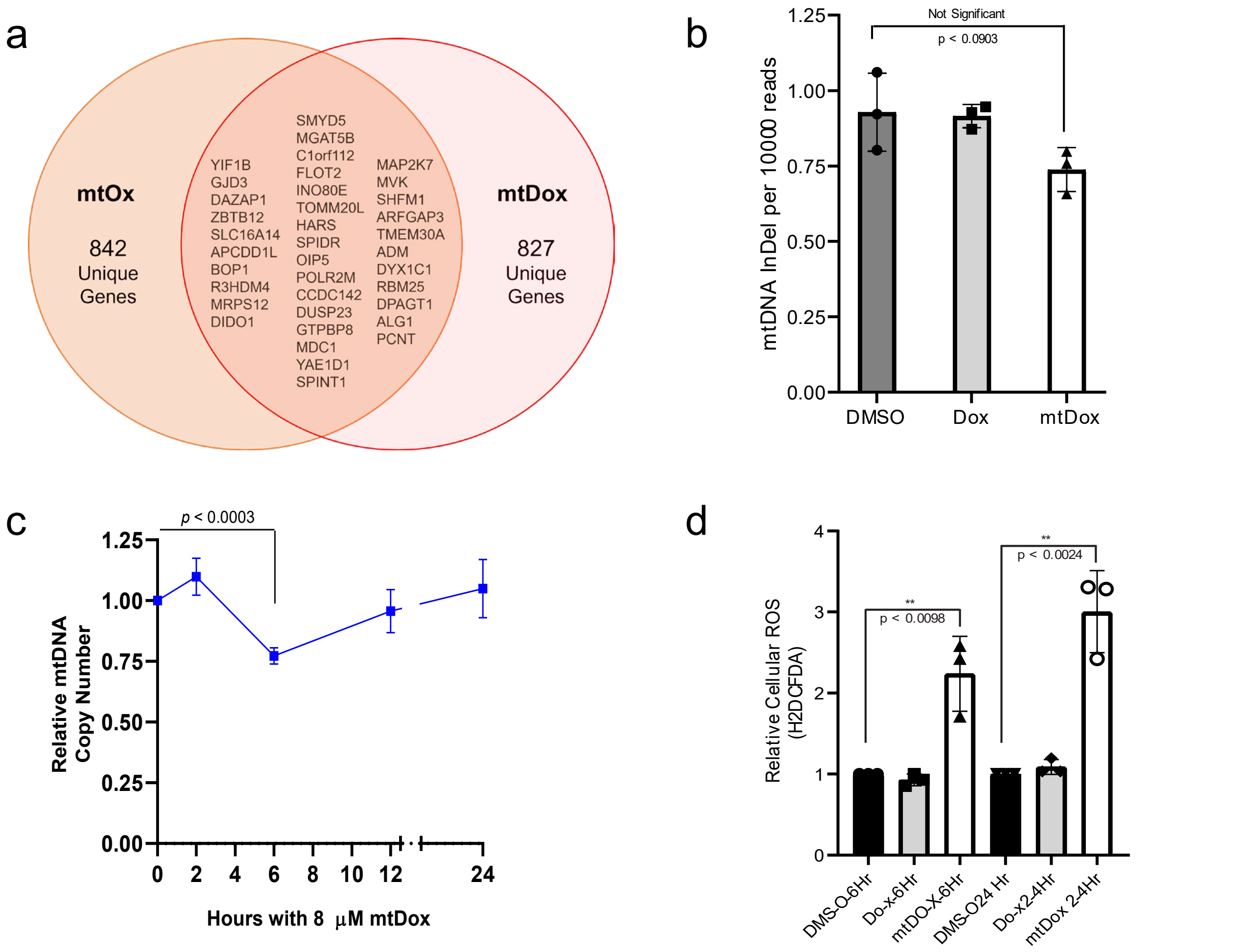


**Figure S4: General effects of mtDox on HCT116 *TP53*(-/-) cells**. **(a)** Demonstration of overlap between different MPP compounds showing the specificity of each conjugate in producing unique hits in genomic CRISPR screens. Compared all genes with a drugZ *p*-value < 0.05 for each compound. **(b)** mtDNA aberration count after 20 days of DMSO, Dox or mtDox treatment. Bowtie was used to compare to consensus mtDNA sequence, then variants were called and counted using freebayes and bcftools (n = 3, $p < 0.0903$ n.s.). The observed slight decrease in mtDox population can be explained by the greater number of mtDox-treated transcripts analyzed. The number of InDels observed was consistent between populations, likely due to pre-existing sequence differences between the reference genome and the analyzed genome. The number of InDels was normalized to mtDNA reads, causing the mtDox population to have relatively less InDels per read analyzed. **(c)** Measurement of mtDNA copy number over time with 8 μM mtDox treatment using qPCR (n = 3, $p < 0.0003$ at 6 hours post-treatment). **(d)** Summarized flow cytometry analysis of total cellular ROS using H2DCFDA probe. Cells were treated for indicated amount of time with Dox or mtDox concentrations of 25 nM Dox and 8 μM mtDox to match conditions used for the CRISPR/Cas9 screen validation experiments (n = 3, $p < 0.0098$ for mtDox at 6 hours, $p < 0.0024$ for mtDox at 24 hours. All *p*-values determined using unpaired *t*-test. Data represented as mean ± s.d.

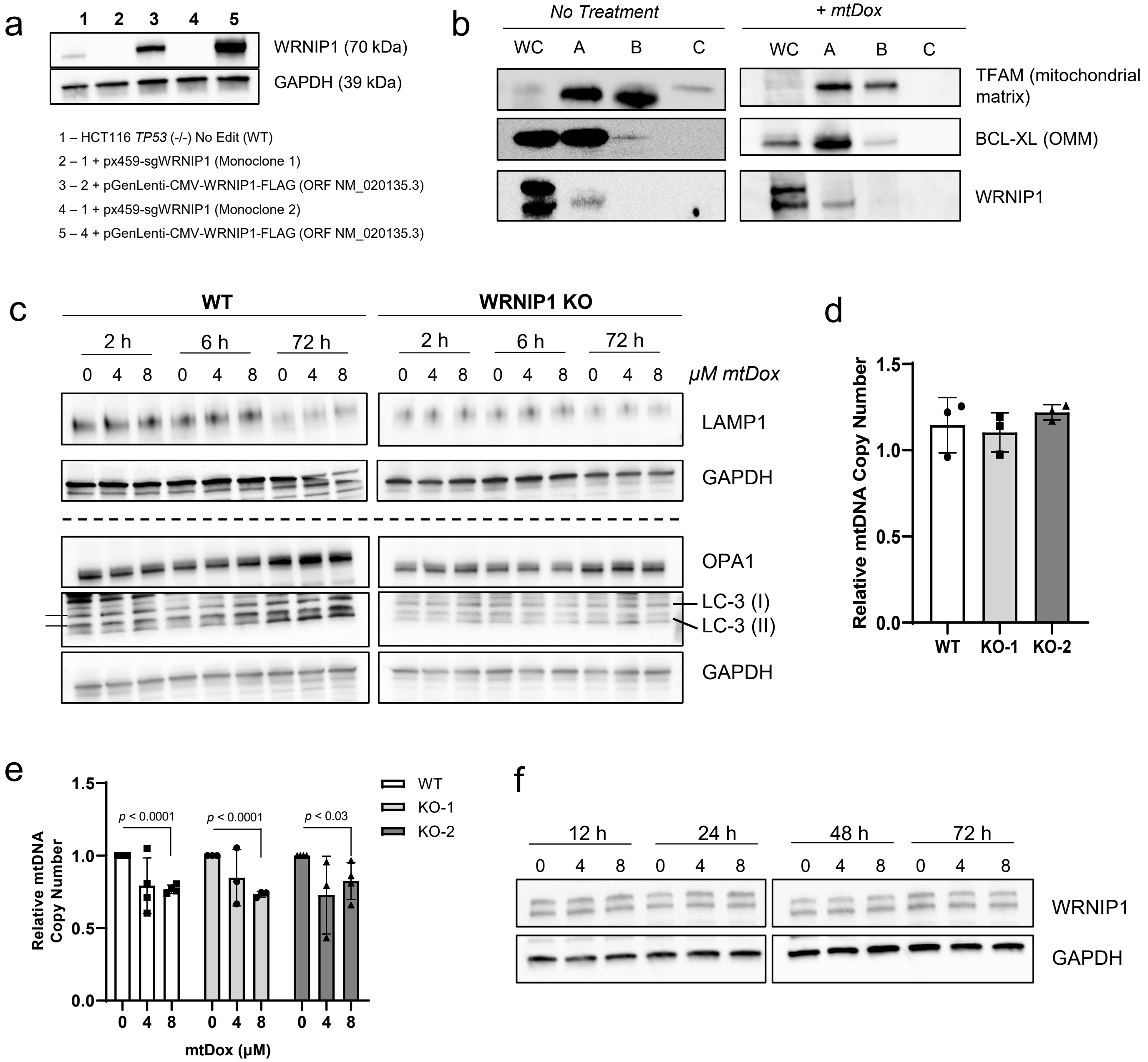


**Figure S5: Additional experiments for mitochondrial role of WRNIP1. (a)** Validation and identity of monoclonal WRNIP1 knockout (KO) lines along with their reintroduction of WRNIP1 through a cDNA construct pGenLenti. **(b)** Further analysis of proteinase K mitochondrial localization test with 24 hour, 4 µM mtDox treatment. WC = whole cell, A = mitochondrial isolation via ultracentrifugation, B = A + 100 µg/mL proteinase K, C = B + 10% SDS. **(c)** Western blot analysis of potential mtDox mediated death pathways and mitochondrial network rewiring. **(d)** Comparing basal mtDNA copy number between WT HCT116 *TP53* (-/-) cells and two WRNIP1 monoclonal knockouts (KO) using qPCR. No significant difference was found between populations (n = 3). **(e)** Comparing trend of 6-hour concentration dependent mtDox treatment between WT cells and two WRNIP1 monoclonal knockouts (KO) using qPCR. All three populations display a similar level of mtDNA copy number reduction from treatment (n = 3). For **d** and **e**, all *p*-values determined using unpaired *t*-test. Data points represented as mean ± s.d. **(f)** Western blot time-course of WRNIP1 expression over time with increasing mtDox treatment concentrations. WRNIP1 expression was found to be unaffected by mtDox.

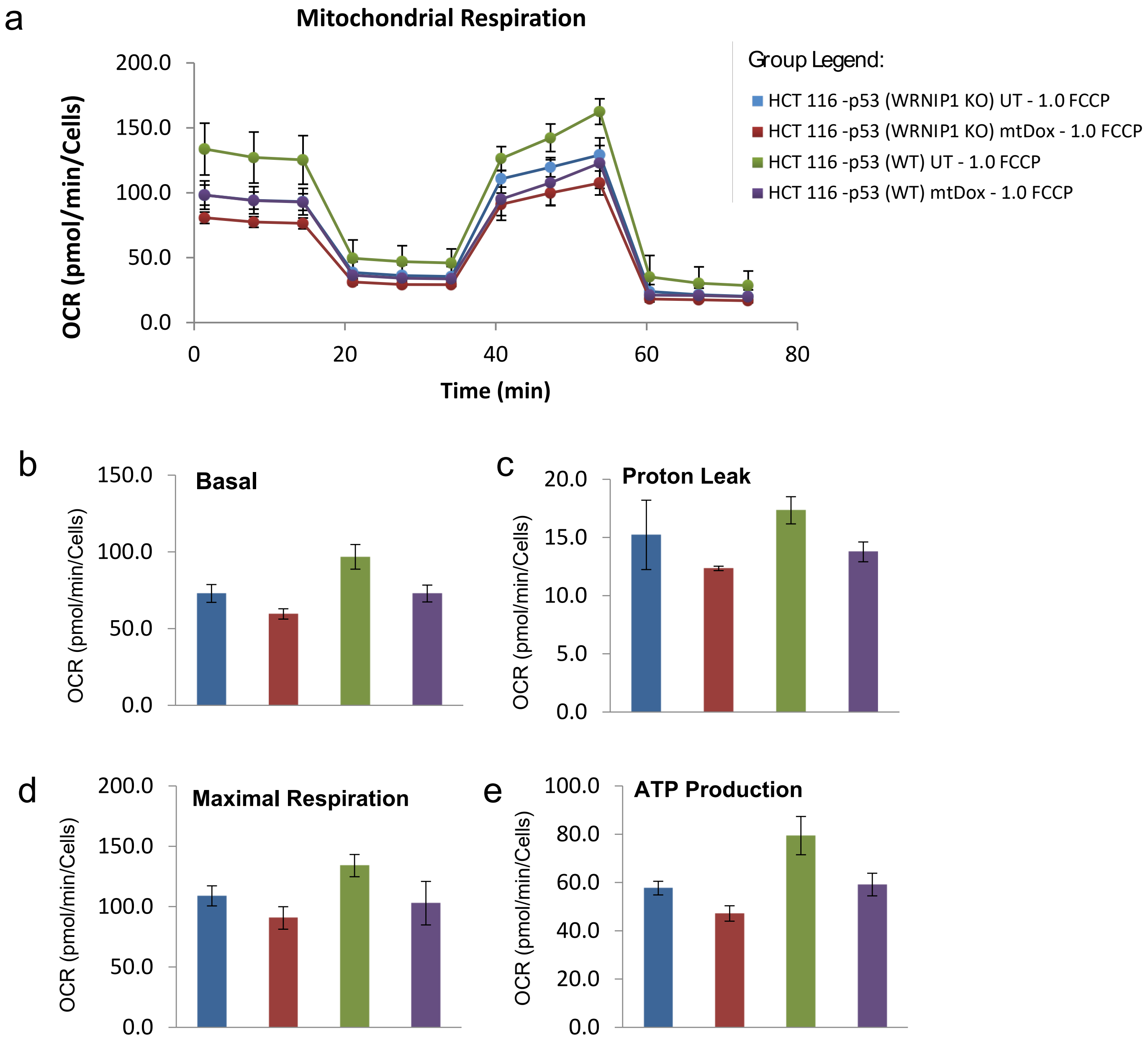


**Figure S6: Seahorse analysis of the effect of WRNIP1 knockout and mtDox treatment. (a)** Full time-course representation of Seahorse Mito Stress Test with 1.0 μM FCCP. **(b - e)** Highlighted data trends from **a** showing differential oxygen consumption rate (OCR) at proceeding stages of the experiment (n = 4).

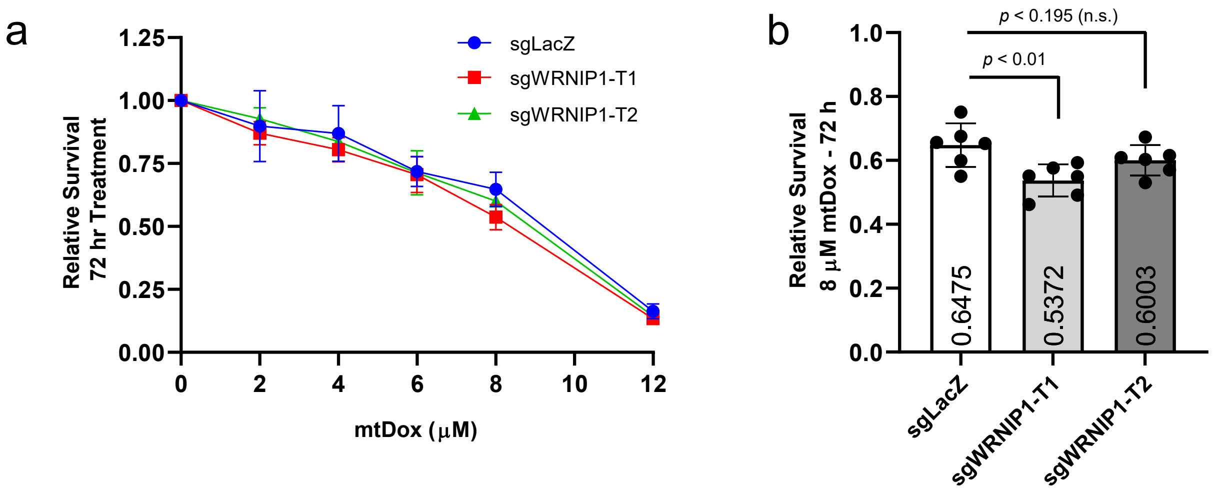


**Figure S7: Analysis of short-term WRNIP1 KO survivability. (a)** Relative survival normalized to untreated cells using PrestoBlue Cell Viability Reagent for sgLacZ, sgWRNIP1-T1, and sgWRNIP1-T2 at 72 hours mtDox treatment. **(b)** Highlighted survivability at 8 μM mtDox for comparison with previous clonogenic survival tests with a duration of 14 days treatment (n = 6). All *p*-values determined using unpaired *t*-test. Data represented as mean ± s.d.

### Source Data – Western Blots

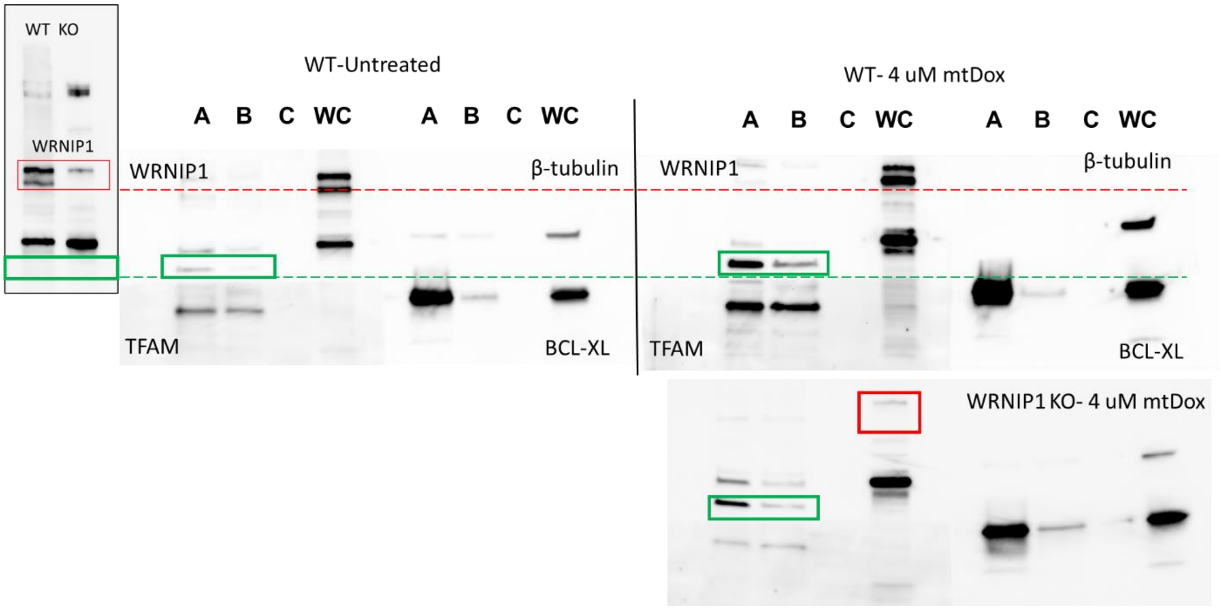


**Source Data – Figure 3b (and Figure S5b) Full Western blot of proteinase K analysis of mitochondrial localization.** WRNIP1 band identity confirmed by knockout (red box and dotted line). Treatment with mtDox did not affect mitochondrial localization. Did see the appearance of a different sized band with mtDox treatment (green box and dotted line), however this band did not disappear with WRNIP1 knockout and could not be confirmed to be related to WRNIP1.

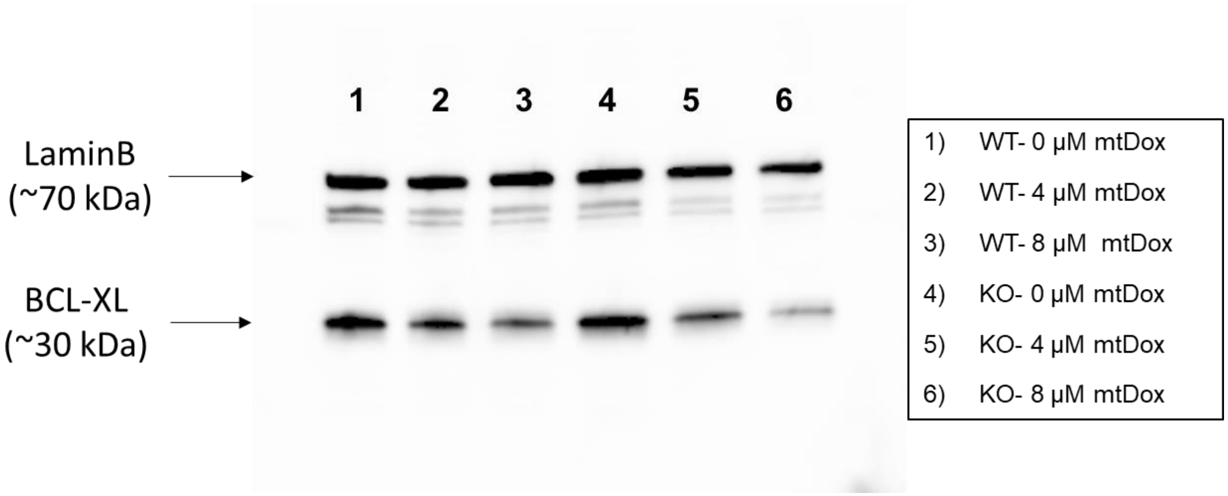


**Source Data – Figure 3g: Full Western blot of BCL-XL expression with LaminB loading control**. Blot was cut to probe for multiple targets from same isolates. Dashed blue line represents where blot was cut to allow probing of both antibodies from same blot.

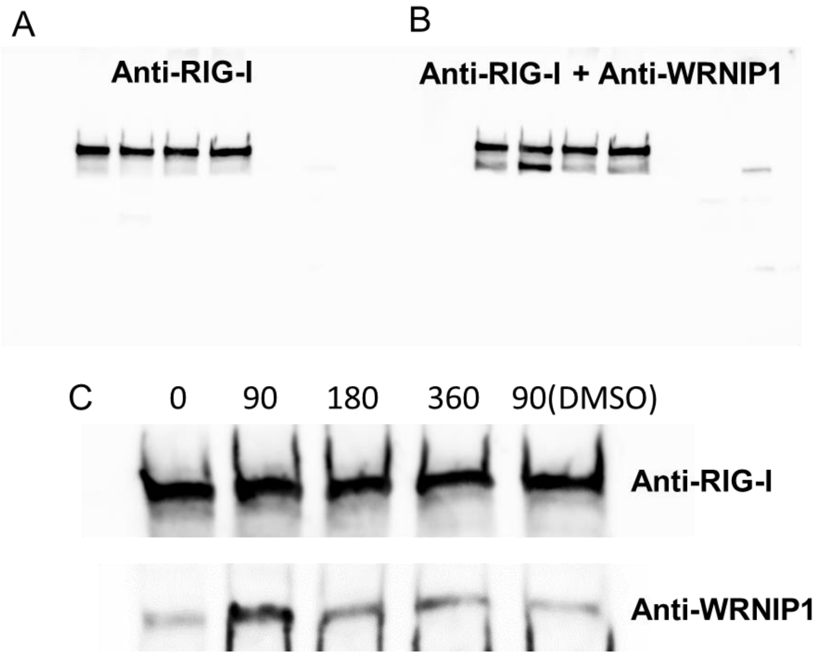


**Source Data- Figure 4d: Full Western blot of RIG-I co-immunoprecipitation.** A, B) Performed sequential staining because bands were too close to cut between. Faint two lanes to the right of the 4 main bands represent anti-WRNIP1 staining of WT versus WRNIP1 KO cells, confirming band size for WRNIP1. C) Another replicate of the co-immunoprecipitation including DMSO control. All other numbers represent minutes of 8 µM mtDox treatment.

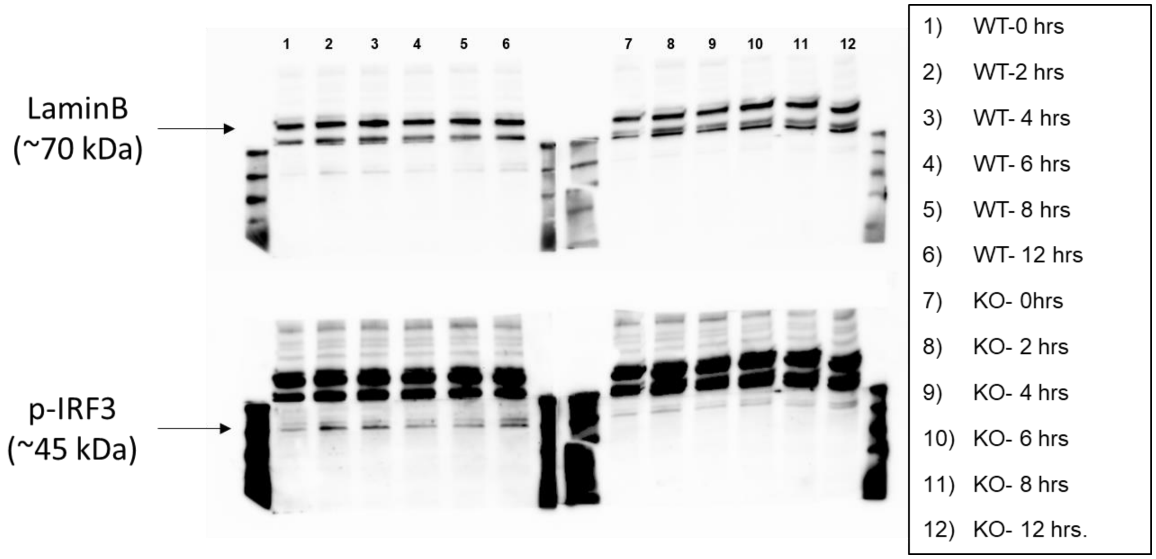


| p-IRF3/LaminB (normalized) | 0hr | 2hr | 4hr | 6hr | 8hr | 12hr |
|---|---|---|---|---|---|---|
| WT | 1 | 4.01 | 3.70 | 2.83 | 2.74 | 5.44 |
| KO | 1 | 1.06 | 0.92 | 0.66 | 0.98 | 2.34 |

**Source Data Figure 4e: Full Western blot of p-IRF3 assay for WT versus WRNIP1 KO cells with exposure to 8 μM mtDox.** Top and bottom blots represent same experiment with different exposures due to the relatively low expression of p-IRF3 and the relatively high expression of LaminB. Blot was cut to probe for multiple targets from same isolates.

### Source Data – Flow Cytometry

#### Source Data for Figure 3d

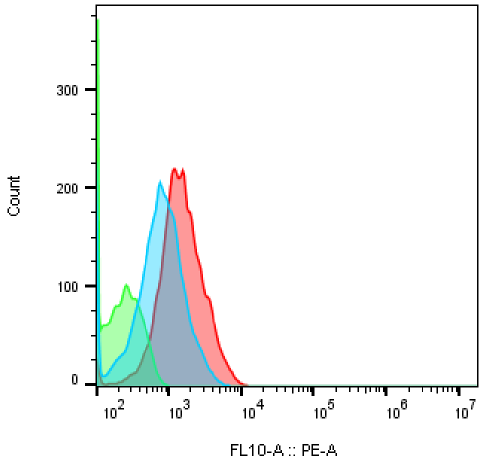


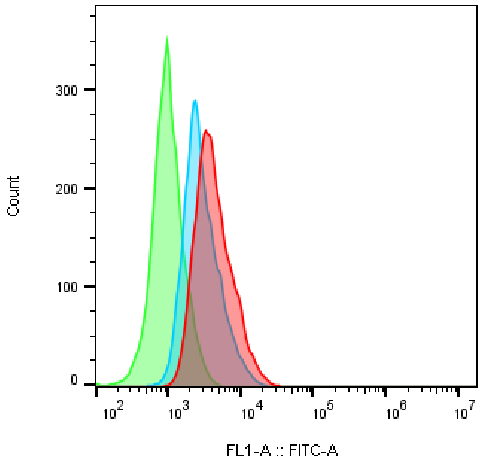


| | Count | Sample Name | Mean : FL10-A |
|---|---|---|---|
| (blue) | 7369 | A7-4uM-Unstained.fcs | 951 |
| (green) | 7804 | A7-DMSO-Unstained.fcs | 174 |
| (red) | 7477 | A7-8uM-Unstained.fcs | 1834 |

| | Count | Sample Name | Mean : FL1-A |
|---|---|---|---|
| (red) | 7477 | A7-8uM-Unstained.fcs | 4978 |
| (blue) | 7369 | A7-4uM-Unstained.fcs | 3399 |
| (green) | 7804 | A7-DMSO-Unstained.fcs | 1103 |

**Control Experiment for JC-1 with mtDox.** mtDox bleeds into both the PE-A and FITC channel therefore did not use any mtDox-treated JC-1 experiments in manuscript.

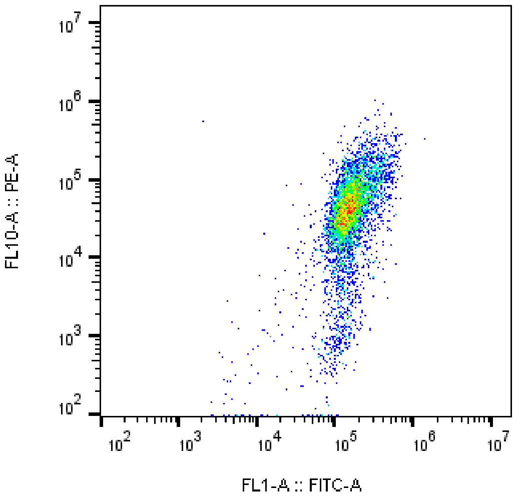


| | Count | Sample Name | Mean : FL10-A | Mean : FL1-A |
|---|---|---|---|---|
| ■ | 5805 | JC-WT-0-1-2.fcs | 57970 | 169026 |

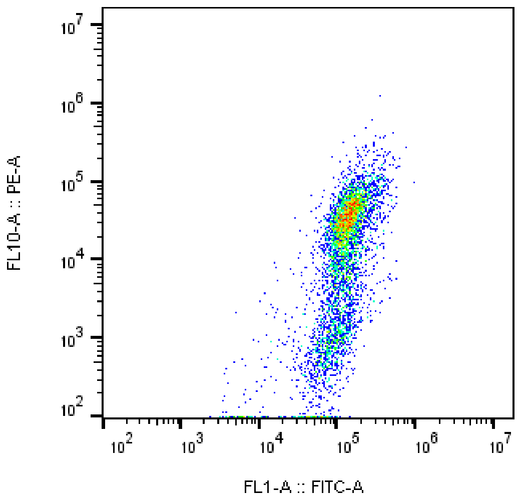


| | Count | Sample Name | Mean : FL10-A | Mean : FL1-A |
|---|---|---|---|---|
| ■ | 4972 | JC-KO-0-1-2.fcs | 29005 | 133163 |

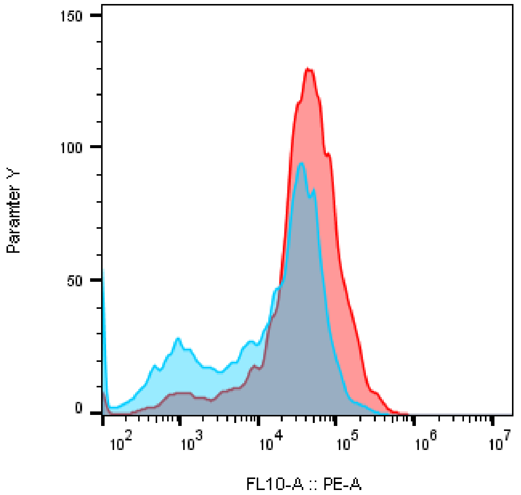


| | Count | Sample Name | Mean : FL10-A | Mean : FL1-A |
|---|---|---|---|---|
| ■ | 4972 | JC-KO-0-1-2.fcs | 29005 | 133163 |
| ■ | 5805 | JC-WT-0-1-2.fcs | 57970 | 169026 |

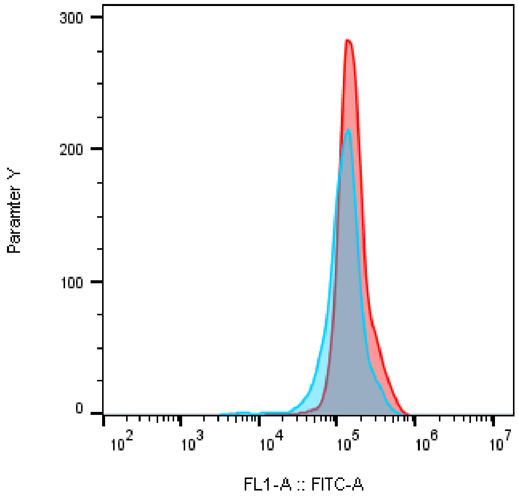


| | Count | Sample Name | Mean : FL10-A | Mean : FL1-A |
|---|---|---|---|---|
| ■ | 4972 | JC-KO-0-1-2.fcs | 29005 | 133163 |
| ■ | 5805 | JC-WT-0-1-2.fcs | 57970 | 169026 |

**(Above) Raw JC-1 experiment data showing reduced basal membrane potential in monoclonal KO-1.**

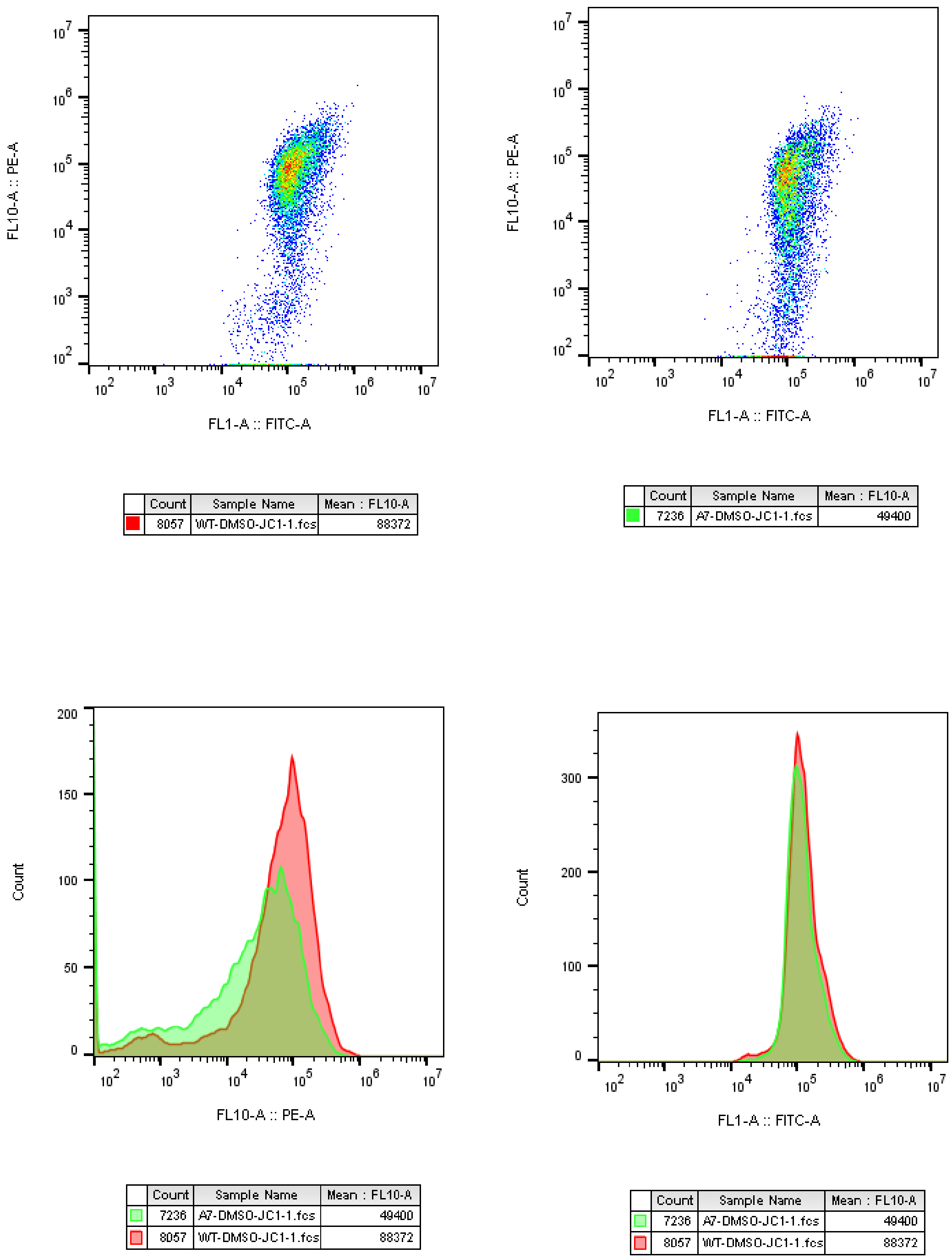


**(Above) Raw JC-1 experiment data showing reduced basal membrane potential in monoclonal KO-2.**

**Source Data-Figure 3 e,f**

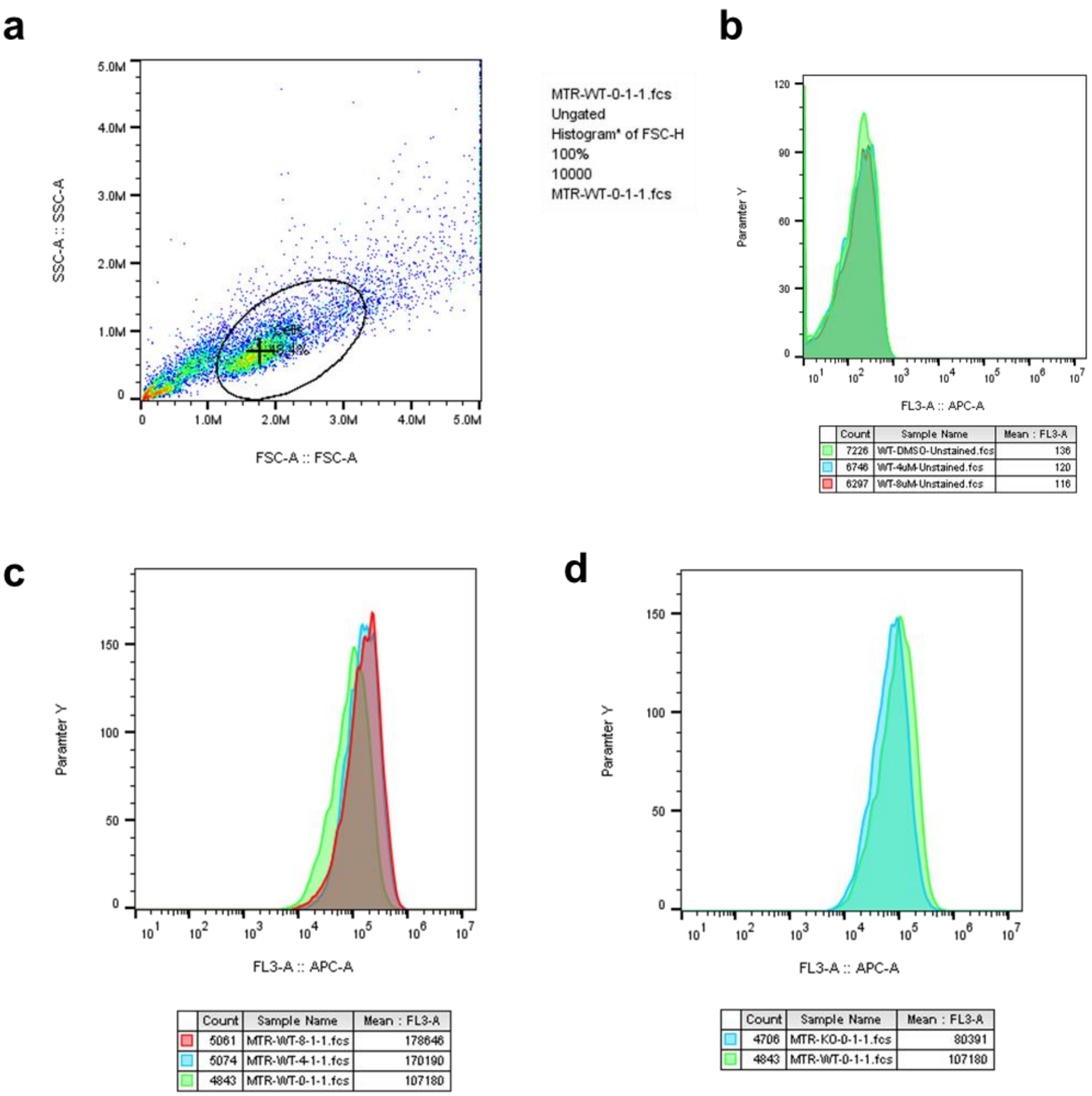


**(Above) Raw data for MitoTracker Deep Red flow cytometry.** a) Gating strategy for cells based on forward and side scattering. b) Control experiment showing no overlap with mtDox and the APC channel used for analysis. c) Comparative data between 0, 4, and 8 μM mtDox. d) Comparative data for WT vs WRNIP1 KO.

#### Source Data-Extended Data Fig. 4c:

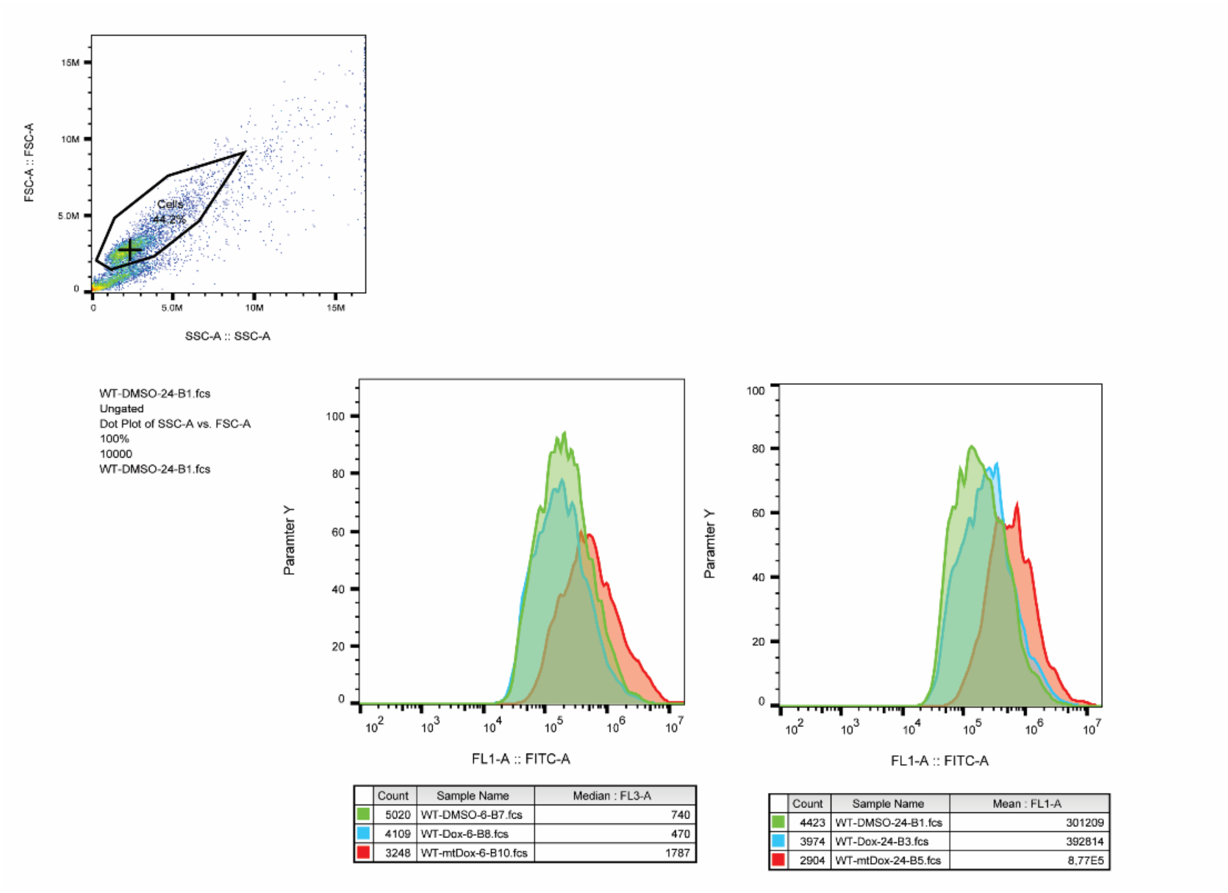


**(Above) H2DCFDA analysis raw data with gating strategy.**

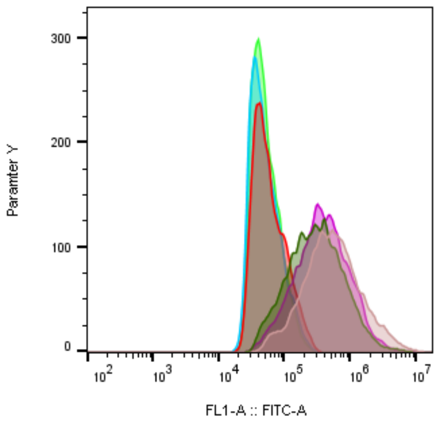


| | Count | Sample Name | Median : FL1-A | Sample Name | Mean : FL1-A |
|---|---|---|---|---|---|
| | 6234 | WT mtDOX DF.fcs | 4,94E5 | WT mtDOX DF.fcs | 8,67E5 |
| | 6824 | WT Dox DF.fcs | 287118 | WT Dox DF.fcs | 4,94E5 |
| | 6922 | WT DF Untreated.fcs | 358925 | WT DF Untreated.fcs | 5,74E5 |
| | 6342 | WT mtDOX Unstained.fcs | 51355 | WT mtDOX Unstained.fcs | 67274 |
| | 6886 | WT Dox Unstained.fcs | 44733 | WT Dox Unstained.fcs | 57408 |
| | 7037 | WT Unstained.fcs | 46066 | WT Unstained.fcs | 57498 |

**(Above) H2DCFDA with unstained controls showing the lack of overlap with mtDox.**
